# Biochemical, biophysical, and structural investigations of two mutants (C154Y and R312H) of the human Kir2.1 channel involved in the Andersen-Tawil syndrome

**DOI:** 10.1101/2024.02.09.579451

**Authors:** Dania Zuniga, Andreas Zoumpoulakis, Rafael F. Veloso, Laurie Peverini, Sophie Shi, Alexandre Pozza, Valérie Kugler, Françoise Bonneté, Tahar Bouceba, Renaud Wagner, Pierre-Jean Corringer, Carlos A. H. Fernandes, Catherine Vénien-Bryan

## Abstract

Inwardly rectifying potassium (Kir) channels play a pivotal role in physiology by establishing, maintaining, and regulating the resting membrane potential of the cells, particularly contributing to the cellular repolarization of many excitable cells. Dysfunction in Kir2.1 channels is implicated in several chronic and debilitating human diseases for which there are currently no effective treatments. Specifically, Kir2.1-R312H and Kir2.1-C154Y mutations are associated with Andersen-Tawil syndrome (ATS) in humans. We have investigated the impact of these two mutants in the trafficking of the channel to the cell membrane and function in *Xenopus laevis* oocytes. Despite both mutations being successfully trafficked to the cell membrane and capable of binding PIP_2_ (phosphatidylinositol-4,5- bisphosphate), the main modulator for channel activity, they resulted in defective channels that do not display K^+^ current, albeit through different molecular mechanisms. Co-expression studies showed that R312H and C154Y are expressed and associated with the WT subunits. While WT subunits could rescue R312H dysfunction, the presence of a unique C154Y subunit disrupts the function of the entire complex, which is a typical feature of mutations with a dominant-negative effect. Molecular dynamics simulations showed that Kir2.1-C154Y mutation induces a loss in the structural plasticity of the selectivity filter, impairing the K^+^ flow. In addition, the cryo-EM structure of the Kir2.1-R312H mutant has been reconstructed. This study identified the molecular mechanisms by which two ATS-causing mutations impact Kir2.1 channel function and provide valuable insights that can guide potential strategies for the development of future therapeutic interventions for ATS.

## Introduction

Potassium channels are a very large family of integral membrane proteins that are ubiquitous with incredibly diverse physiological functions. There are more than 80 genes that code for K^+^ channels in the human genome alone (1). The differences in their structures allow them to be regulated by various auxiliary subunits, to interact with different agonists, and to intervene at different stages of the action potential with diverse effects. Kir channels were first described in frog skeletal muscle as having an anomalous rectification (2) because their conductance decreased instead of increasing upon cell depolarization. Although outward Kir currents are much smaller than inward ones, under physiological conditions, the outward currents regulate the excitability and duration of repolarization in excitable cells like neurons and cardiac myocytes (3). Kir channels are responsible for generating and propagating the action potential in excitable cells, regulating the heartbeat, maintaining salts and water balance, and regulating cell volume and muscle contraction. To fulfill these crucial biological roles, the gating of all eukaryotic Kir channels is directly controlled by a variety of intracellular messengers or regulatory lipids (3) The Kir channels family is classified into seven subfamilies (Kir1 to Kir7) with Kir2.1 being the protein product of the KCNJ2 gene (3). Kir2.1 is expressed at high levels in the heart, skeletal muscle, and neural tissue, and Kir2.1 mutations are associated with various diseases (4). Kir2.1 channels are selectively modulated by the signaling lipid phosphatidylinositol 4,5- bisphosphate (PIP_2_) (5). Loss-of-function mutations in Kir2.1 channel cause Andersen-Tawil syndrome (ATS), a rare autosomal disorder characterized by developmental skeletal abnormalities, periodic skeletal muscle paralysis, as well as biventricular tachycardia with or without the presence of long QT (6, 7). On the other hand, gain-of-function mutations in Kir2.1 result in cardiac phenotypes, including atrial fibrillation, and short QT syndrome (SQT3), attributed to increased repolarization capacity and thus shortened cardiac action potentials (8, 9).

All Kir channels are tetrameric structures and are thought to assemble onto tetramers of identical or related subunits in the endoplasmic reticulum (ER) of cells (10). Kir channels control the passage of K^+^ ions through the selectivity filter containing the conserved amino acids ^142^TIGYG^146^ (Kir2.1 numbering) (11). Kir channels share characteristic structural features: a large cytoplasmic domain (CTD) hosting the C- and N-termini and a transmembrane domain (TMD) containing a slide helix, a pore helix, a loop containing the K^+^ ion selectivity filter and two transmembrane helices (M1 and M2) separated by an extracellular loop (12, 13). In the Kir2.1 structure, the tetrameric assembly is mediated mainly through interactions involving the M2, selectivity filter, and pore helix at the TMD, as well as the G-loop (300-315 region), and regions 226-250, and 325-340 at the CTD. No inter-subunit disulfide bonds are involved in the tetrameric assembly (12).

The Kir2.1-R312H and Kir2.1-C154Y mutants were first identified in 2009 and have been reported as loss-of-function mutations responsible for ATS in humans (14). The residue R312 is located in the CTD (Figure 1A), near the G-loop and the PIP_2_ putative binding site, but is not directly involved in the PIP_2_ binding. Nevertheless, this mutant does not conduct K^+^. In our previous study, this residue has been described as important for the signal transmission to the G-loop gating mechanism upon PIP_2_ binding (12). In contrast, the residue C154 is located in the N-terminal of the M2 helix and forms the unique disulfide bond within the Kir2.1 structure with the C122 of the same chain (Figure 1A-B). The residues C122 and C154 are conserved in all eukaryotic Kir members (Figure 1C). The mechanism by which the C154Y mutation renders the channel deficient remains controversial. Previous studies have suggested that the C122/C154 disulfide bond in Kir2.1 appears to be necessary for the channel assembly, but less critical for function in an assembled channel (15, 16). Recently, deep mutational scanning (DMS) experiments on mouse Kir2.1 expressed in HEK293T cells have indicated that mutations in the C154 residue may possibly disrupt cell surface expression to a certain extent but not function (17). On the other hand, the crystal structure of chicken Kir2.2 has suggested that this disulfide bond may indeed be significant for channel function (18). Moreover, studies on Kir3.2 channels have demonstrated that these cysteine residues are not essential for the correct processing and insertion of the channel into the membrane but are crucial for its function (19). Interinstingly, all prior experiments performed in *Xenopus laevis* oocytes have shown that mutated C154Y channels are expressed at the membrane (15, 16).

**Figure 1.**
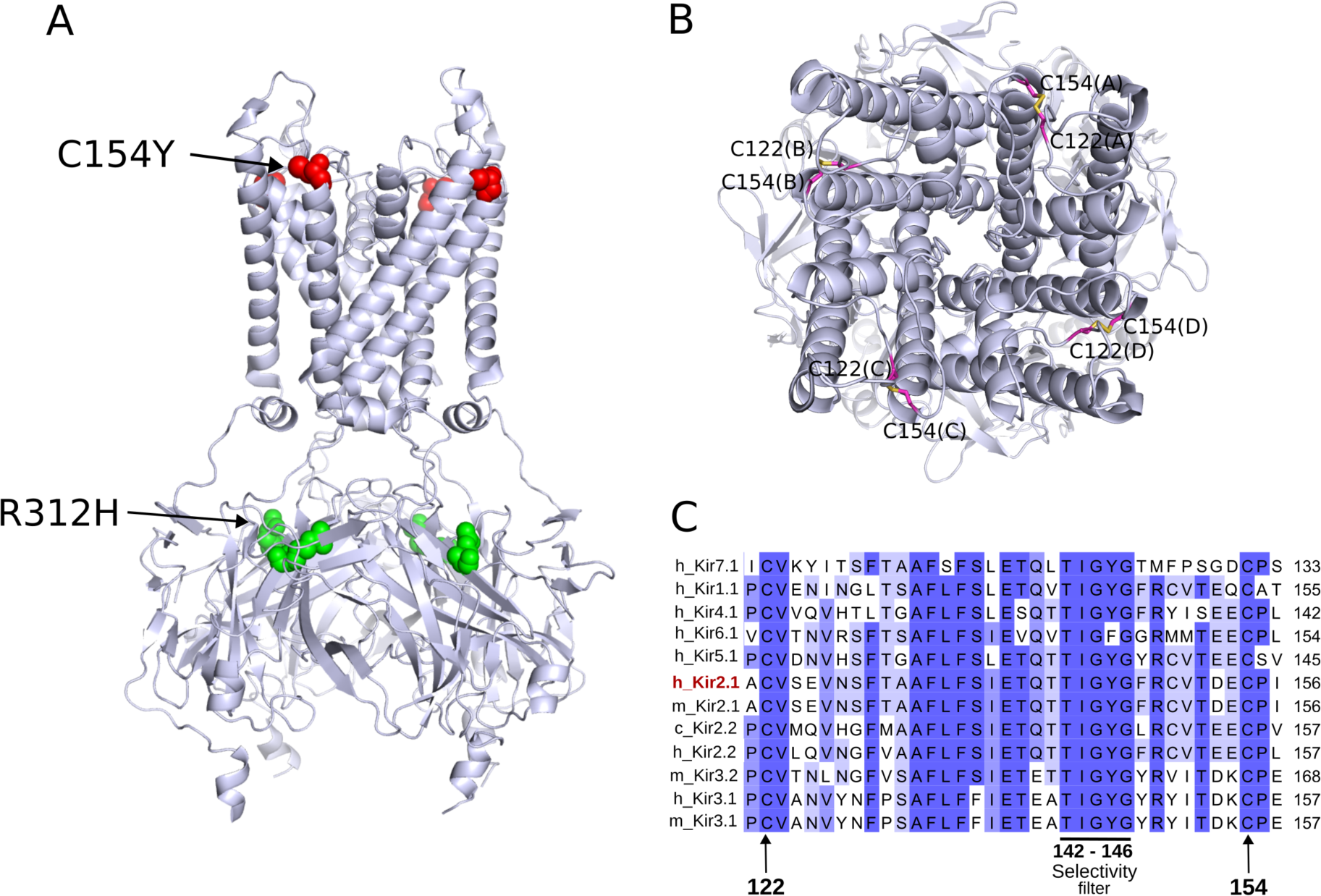
**Location of C154Y and R312H mutations associated with Andersen-Tawil syndrome on human Kir2.1 channel. A**. Cartoon representation of the cryo-EM structure of the human Kir2.1 channel (PDB ID 7ZDZ) where the location of the C154Y and R312H mutations is highlighted on the channel structure (red and green spheres, respectively). **B**. Top view of the human Kir2.1 channel structure where the intra-subunit disulfide bond formed by C122 and C154 is highlighted in magenta sticks. The chains are indicated in parentheses. **C**. Amino acid sequence alignment of the region comprising residues 121 to 155 (Kir2.1 numbering) between different Kir channels. The conserved C122 and C154 residues are indicated by black arrows (numbers 122 and 154). The alignment also highlights the selectivity filter of Kir channels (residues 142- 146). The highly conserved amino acids are highlighted in purple. h_Kir: Human Kir, m_Kir: mouse Kir, c_Kir: chicken Kir. The amino acid sequence of the human Kir2.1 channel (h_Kir2.1) is highlighted in red.

Herein, we highlighted by different approaches that Kir2.1-C154Y and Kir2.1-R312H mutations do not impact channel trafficking to the membrane on *X. laevis* oocytes. Moreover, we have investigated the effects of these mutations on the structure and function of Kir2.1 channels by integrating results from electrophysiology experiments and structural data obtained through cryo-electron microscopy (cryo-EM) single-particle analysis and molecular dynamics (MD) simulations. In this paper, we propose two distinct molecular mechanisms associated with each studied mutation, both leading to the same pathology, ATS. The results presented here offer valuable insights into the intricate relationship between protein dysfunction and disease manifestation.

## Results

### Functional properties of Kir2.1-WT and mutants R312H and C154Y in *Xenopus* **oocytes.**

We studied the effects of replacing Arg312 with His (R312H) and Cys154 with Tyr (C154Y) on the K^+^ current of Kir2.1 channels by recording two-electrode voltage clamp (TEVC) currents in *Xenopus laevis* oocytes (Figure 2). Upon injection of 50 ng of cDNA of the Kir2.1-WT, the oocytes showed classical currents with strong rectification properties (Figure 2, in dark blue). The maximum negative current at -150 mV (-Imax) registered for oocytes injected with 50 ng of Kir2.1-WT cDNA was normalized to 100% activity in this study. (Figure 2 in dark blue). The oocytes injected with 50 ng of cDNA of the mutants R312H and C154Y did not display observable currents and were indistinguishable from endogenous currents recorded from non-injected oocytes (Figure 2, in green and red on panels A and B, respectively). To investigate whether the Kir2.1-R312H and Kir2.1-C154Y mutants are able to associate with WT subunits and form functional channels, we co-expressed Kir2.1-WT cDNA together with either mutant R312H or C154Y cDNAs (Figure 2). Below, we make all the comparisons for the various expression relative to 25 ng wild type (Figure 2 blue), the amount of WT present in all mixed cDNA injections and we are discussing in terms of how co-expressing the mutant and WT affects the baseline function of 25 ng WT. The coexpression of R312H mutant mixed 1:1 with WT cDNA (25 ng of Kir2.1-WT cDNA with 25 ng of the Kir2.1- R312H cDNA) shows more activity than 25 ng WT (Figure 2A, in cyan), about 25% increase. An increase in activity over WT values indicates that a greater number of functional channels are present in the population. This enhancement must represent the summation of currents from the WT and functional heteromeric channel, indicating that at certain ratios, WT subunits can rescue defective R312H channel subunits, as macroscopic currents of the mutants alone cannot be measured. Additionally, these data indicate that R312H does not affect the association of subunits to form tetrameric channels.

**Figure 2.**
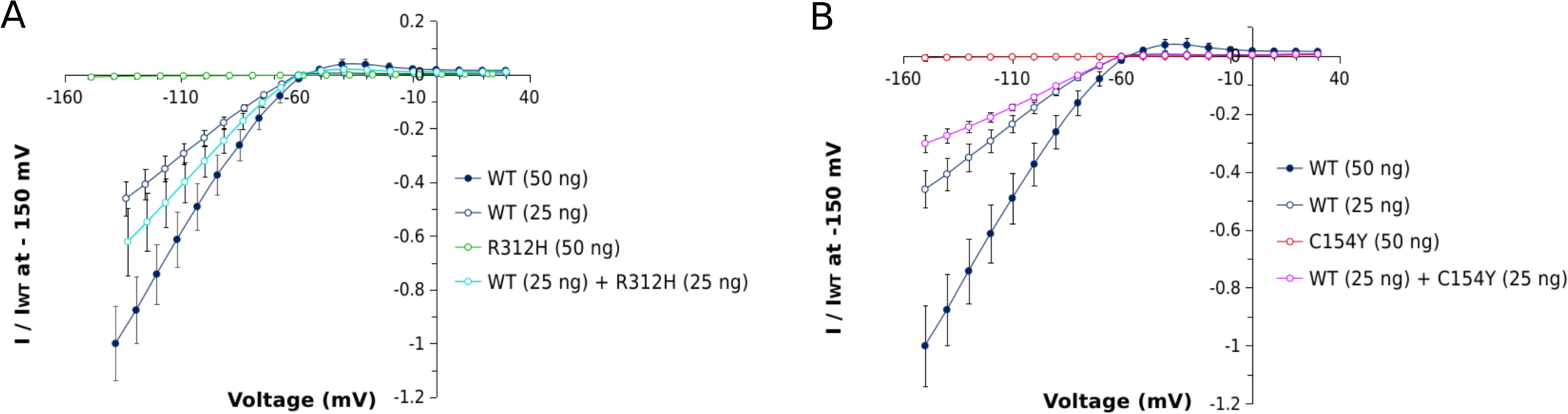
Functional expression of Kir2.1-WT and C154Y and R312H mutants in *Xenopus laevis* oocytes. Normalized current-voltage (I-V) relationships of the macroscopic currents of the Kir2.1 WT alone (injections of 50 ng and 25 ng of WT cDNA, blue and gray curves, respectively) and (**A**) R312H mutant alone (50 ng of R312H cDNA, green curve) or co-injected with WT cDNA (25 ng of WT cDNA + 25 ng of R312H cDNA, cyan curve), and (**B**) C154Y mutant alone (50 ng of C154Y cDNA, red curve) or co-injected with WT cDNA (25 ng of WT cDNA + 25 ng of C154Y cDNA, purple curve) compared to the control current of Kir2.1-WT (50 ng and 25 ng of cDNA, blue and gray curves) at −150 mV. Currents were elicited by 500 ms pulses from −150 mV to +30 mV in 10 mV increments from a holding potential of −60 mV. Values were measured at the end of each step pulse and represent mean ± SEM, n = 6-16 oocytes per group.

In contrast, the 1:1 mixture of C154Y mutant with WT cDNA (25 ng of Kir2.1-WT cDNA with 25 ng of the Kir2.1-C154Y cDNA) shows less activity than 25 ng WT alone, about 30% decrease (Figure 2B, in magenta). This implies that heteromerization with C154Y removes functional channels from the population, as at least some heteromers are non-functional channels.

Unpaired two-sided student t-test indicates the significance of the effect of the mutations on - Imax at -150 mV (Figure S1).

### Immunofluorescence experiments in *Xenopus* oocytes

To determine if the WT, R312H, and C154Y subunits of the Kir2.1 channel are transported to the cell membrane, or if they are retained within the cell, we performed immunofluorescence experiments in *Xenopus laevis* oocytes. Images were obtained with an epifluorescence microscope using an anti-Kir antibody (Figure 3A). The top four horizontal panels highlight the visualization of the orange-fluorescent Cyanine3 (Cy3) dye conjugated to the anti-Kir antibody which shows the location of Kir2.1. Compared to the negative control of GFP-only injected oocytes (top left image of the panel), expression of Kir2.1-WT and R312H and C154Y mutants can be visualized in the membrane surface of the oocytes. The lower four panels show the superimposition of two colors: one emitted from the Cy3 conjugate, and the other emitted from the co-injected green fluorescent protein (GFP), which is used to confirm the successful injection of cDNA into the oocytes. The results show that the WT and mutant forms of the channel are expressed in the cell membrane of *Xenopus* oocytes, indicating that the mutations R312H and C154Y do not significantly impact the trafficking of the protein to the membrane in oocytes.

**Figure 3.**
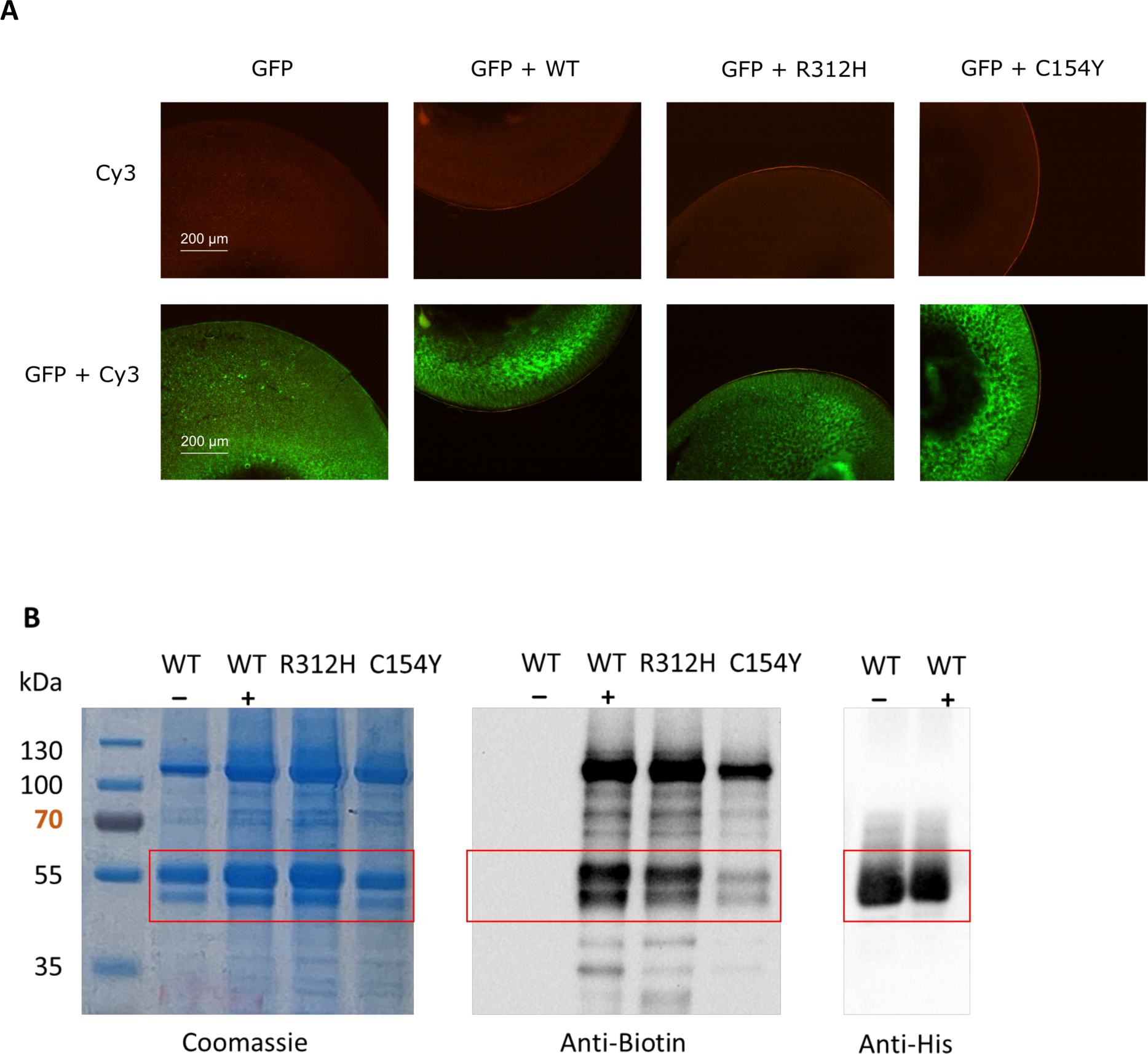
Trafficking of the Kir2.1-WT and R312H and C154Y mutants to the membrane. A. Immunofluorescence assays conducted in *Xenopus laevis* oocytes expressing Kir2.1-WT and mutants. Top row: visualization of Cy3-conjugated anti-rabbit secondary antibody (the primary antibody is a rabbit anti-Kir2.1) showing the location of Kir2.1-WT, Kir2.1-R312H, and Kir2.1- C154Y in the membrane (orange). Control (no cDNA Kir2.1 injected) is shown top left. Bottom row: same as top row but with the co-visualization of the co-injected green fluorescent protein (GFP), which is used to confirm the successful injection of cDNA into the oocytes. Oocytes were co-injected with 50 ng of cDNA encoding Kir2.1 (WT, R312H or C154Y) and 25 ng of cDNA encoding GFP or injected only with 25 ng of cDNA encoding GFP when indicated. Three independent experiments in three different oocytes batches were performed, and the results were consistently similar. One representative experiment is shown in the figure. **B.** Cell surface protein biotinylation assays in *Pichia pastoris* cells. Left: SDS-PAGE of Kir2.1-WT and mutants purified after biotinylation and affinity chromatography revealed by Coomassie blue. The first lane refers to molecular weight markers. Middle: Western blot and Extravidin-HRP reaction highlighting the presence of protein Kir2.1-WT, and R312H and C154Y mutants at the membrane level. WT (-) is the result with the non-biotinylated membrane. Right: Western-blot revealed by Anti-His antibody interaction with His-tagged Kir2.1.

### Expression and purification of the Kir2.1-WT and R312H and C154Y mutant forms

To obtain functional and structural information and perform biochemical and biophysical studies *in vitro*, Kir2.1-WT and both mutant forms were expressed in *Pichia pastori*s cells and purified following the protocol previously described (12). The Kir2.1-R312H mutant exhibited the same level of expression compared with Kir2.1-WT (data not shown) and showed the same size-exclusion chromatography (SEC) profile (Figure S2A). Although the Kir2.1-C154Y mutant did not affect the growth of *P. pastoris* cells, the protein yield was lower than the Kir2.1-R312H and Kir2.1-WT (less than 50%, data not shown). The SEC profile of the Kir2.1-C154Y mutant revealed a peak at the elution volume corresponding to aggregates and a small peak eluting at the same elution volume as the Kir2.1-WT tetramer (Figure S2A, in cyan). This small peak was used to conduct a native PAGE gel, which revealed that this fraction was polydisperse and consisted of aggregates and a faint quantity of tetramers (Figure S2B) in the absence of reducing and denaturing agents. Kir2.1-C154Y mutant was not purified in sufficient quantity to perform SEC-MALLS-RI and cryo-EM analysis but enough for the SPR experiments.

### Biotinylation assays: expression of the Kir2.1-WT and mutants on the cell surface

To verify that the Kir2.1-R312H and Kir2.1-C154Y mutants are trafficked to the plasma membrane in yeast, another expression system for purified hKir2.1, we evaluated the cell surface protein expression by biotinylation assays on whole *Pichia pastoris* cells (Figure 3B). After using a non-permeant reactant biotinylating primary amines at the cell surface, whole membrane proteins were further solubilized, purified, and analyzed using a horseradish peroxidase (HRP) -coupled extravidin probe. While the three Kir2.1-WT, Kir2.1-R312H, and Kir2.1-C154Y constructs were effectively produced and purified from the corresponding expressing clones (Coomassie blue stained SDS-PAGE, Figure 3B, left panel), all the samples submitted to the biotinylation assay were positively detected with extravidin-HRP reactant (Figure 3B, center panel), thus attesting on their presence at the surface of the treated cells These data support our previously obtained immunofluorescence results in *Xenopus* oocytes.

### In solution biophysical characterization of the Kir2.1-WT and R312H mutant by SEC-MALLS-RI

#### Kir2.1-WT solubilized in DDM

Size-exclusion chromatography coupled with multiple angle laser light scattering and differential refractive index (SEC-MALLS-RI) is a suitable technique to determine the absolute mass of the membrane protein-detergent complex, the respective mass fraction of protein, and detergent without the need for column calibration. Additionally, this technique allows the determination of the oligomeric state of the protein (20). The SEC-MALLS-RI profile of the Kir2.1-WT shows a peak compatible with a tetramer at an elution volume of approximately 9.7 mL (Figure 4A). A closer examination of this peak showed that all signals (UV, LS, and ΔRI) aligned perfectly at its center (Figure 4B), allowing determination of the molar mass of the complex (for theoretical details, see supplementary data). In addition, protein conjugate analysis showed that Kir2.1 is extremely homogeneous throughout the peak in the buffer condition used, which includes 0.03% DDM (Figure 4B). The Kir2.1-DDM complex was determined to have a molar mass of 304.9 ± 0.9 kDa, with Kir2.1 contributing 203.9 ± 0.6 kDa and being surrounded by 100.1 ± 0.9 kDa of DDM. This corresponds to a tetrameric protein (theoretical Mw of monomer = 51.6 kDa) with approximately 194 ± 2 bound DDM molecules. The protein-free micelles that eluted at 14.1 mL had a mass of 76.4 ± 1.7 kDa, consistent with the published DDM micelle mass of 72 kDa and corresponding to an aggregation number (Nag) of 140 (20). The solubilization of Kir2.1-WT in FC14 followed by an exchange in DDM or PCC-Malt detergent was also performed and is described in Supplementary data.

**Figure 4.**
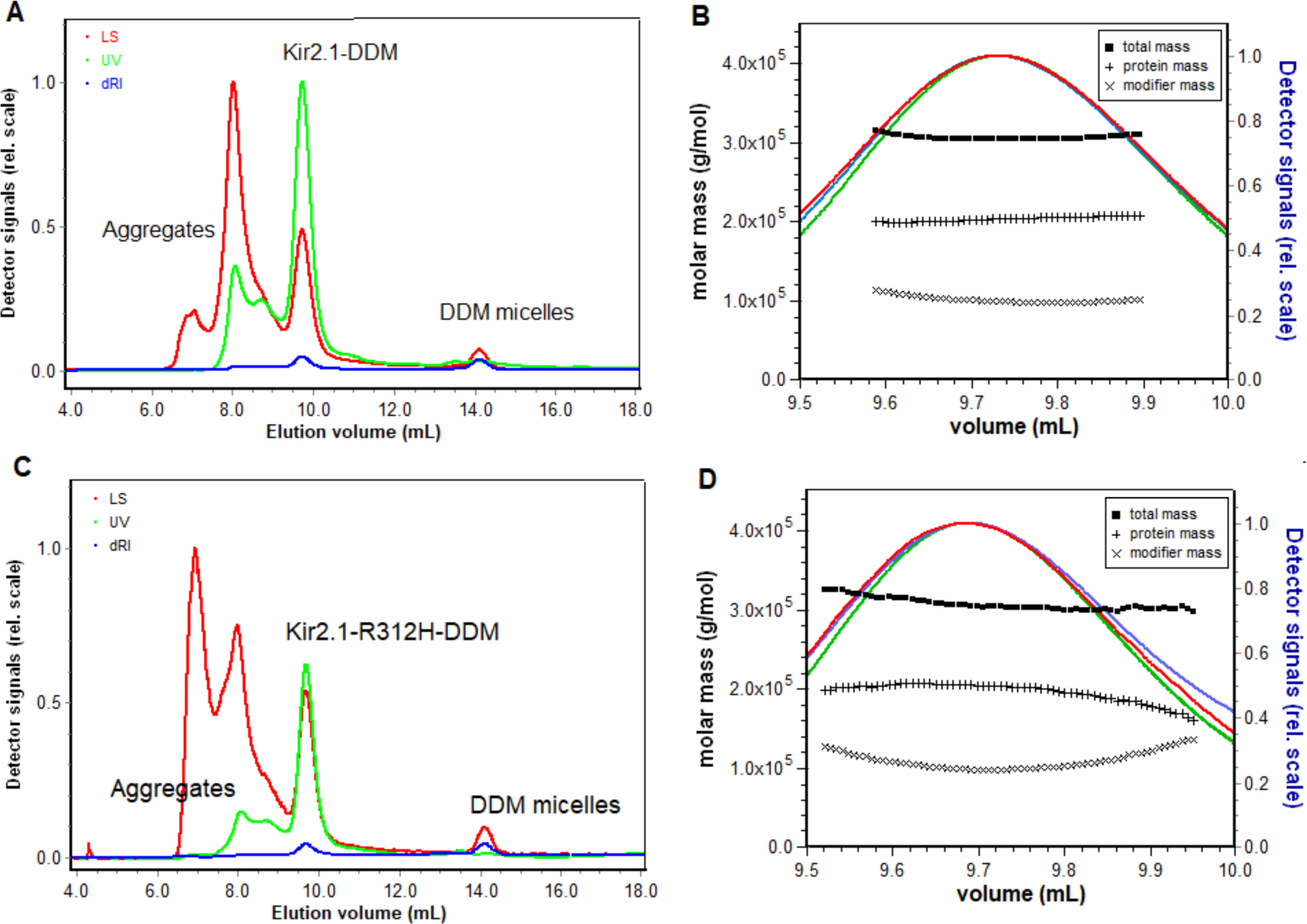
Biophysical characterization of the quaternary structure of Kir2.1-WT and R312H mutant. A. SEC-MALLS-RI profile of Kir2.1-WT in 20 mM Tris-HCl pH 7.4, 150 mM KCl, 1 mM EDTA, 0.03% DDM (eluted on Superdex10/300 200 Increase); **B.** Zoom on the peak corresponding to Kir2.1 tetramers; **C.** SEC-MALLS-RI profile of Kir2.1-R312H in 20 mM Tris-HCl pH 7.4, 150 mM KCl, 1 mM EDTA, 0.03% DDM; **D.** Zoom on the peak corresponding to Kir2.1-R312H tetramers. On each figure, the LS, UV, and RI profiles are shown in red, green, and blue respectively. Molar mass distribution vs elution volume of Kir2.1- DDM and Kir2.1-R312H-DDM complex were calculated using the module Protein Conjugate from the Astra V software (Wyatt Technology). A perfect alignment of the 3 signals (LS, UV, and RI) is observed at the center of the peaks. The total complex mass Kir2.1 molar mass, and DDM molar mass are shown as horizontal lines across the peak, depicting a monodisperse complex in DDM for both Kir2.1-WT and Kir2.1-R312H.

#### Kir2.1-R312H mutant solubilized in DDM

The SEC-MALLS-RI analysis shows that the Kir2.1-R312H-DDM complex was also very homogeneous throughout the peak (Figure 4C-D). This enabled precise calculations of the molar mass of the complex, resulting in a value of 307 ± 4.4 kDa for the complex consisting of tetrameric Kir2.1-R312H with 206.8 ± 3.2 kDa molar mass and 100.3 ± 4.9 kDa DDM. The number of DDM molecules bound to the Kir2.1-R312H-DDM complex was 194 ± 9, similar to that of the Kir2.1-WT-DDM complex (194 ± 2). In conclusion, Kir2.1-R312H formed a homogeneous tetramer, like the Kir2.1-WT counterpart, and therefore, the R312 mutation did not affect the ability of Kir2.1 to form a tetramer.

### PIP_2_ binding interaction with Kir2.1 WT and mutant C154Y

The dissociation constant (K_D_) of Kir2.1 WT and Kir2.1-C154Y mutant to PIP_2_ was determined directly from surface plasmon resonance. Kir2.1-C154Y was immobilized at 3900 to 6500 RU (response units) levels on CM5 sensor chips. Sequential injections of PIP_2_ ranging from 1.25 µM to 20 µM were performed. The fitted curves of the interaction between Kir2.1- C154Y tetramer fraction with PIP_2_ revealed that PIP_2_ was able to bind to the mutant Kir2.1- C154Y (Figure 5) with no significant differences in kinetic parameters between Kir2.1-C154Y (K_D_ = 17.3 µM) and the Kir2.1 WT (K_D_ = 10.3 µM) comparable to the value published in the previous paper (K_D_ = 3.2 ± 0.7 μM) (12) suggesting that the C122/C154Y intra-subunit disulfide bond does not impact the PIP_2_ binding. Likewise, previously obtained SPR results with the Kir2.1-R312H mutant also showed that the R312H mutation did not significantly affect the PIP_2_ binding (K_D_ = 37 μM) (12).

**Figure 5.**
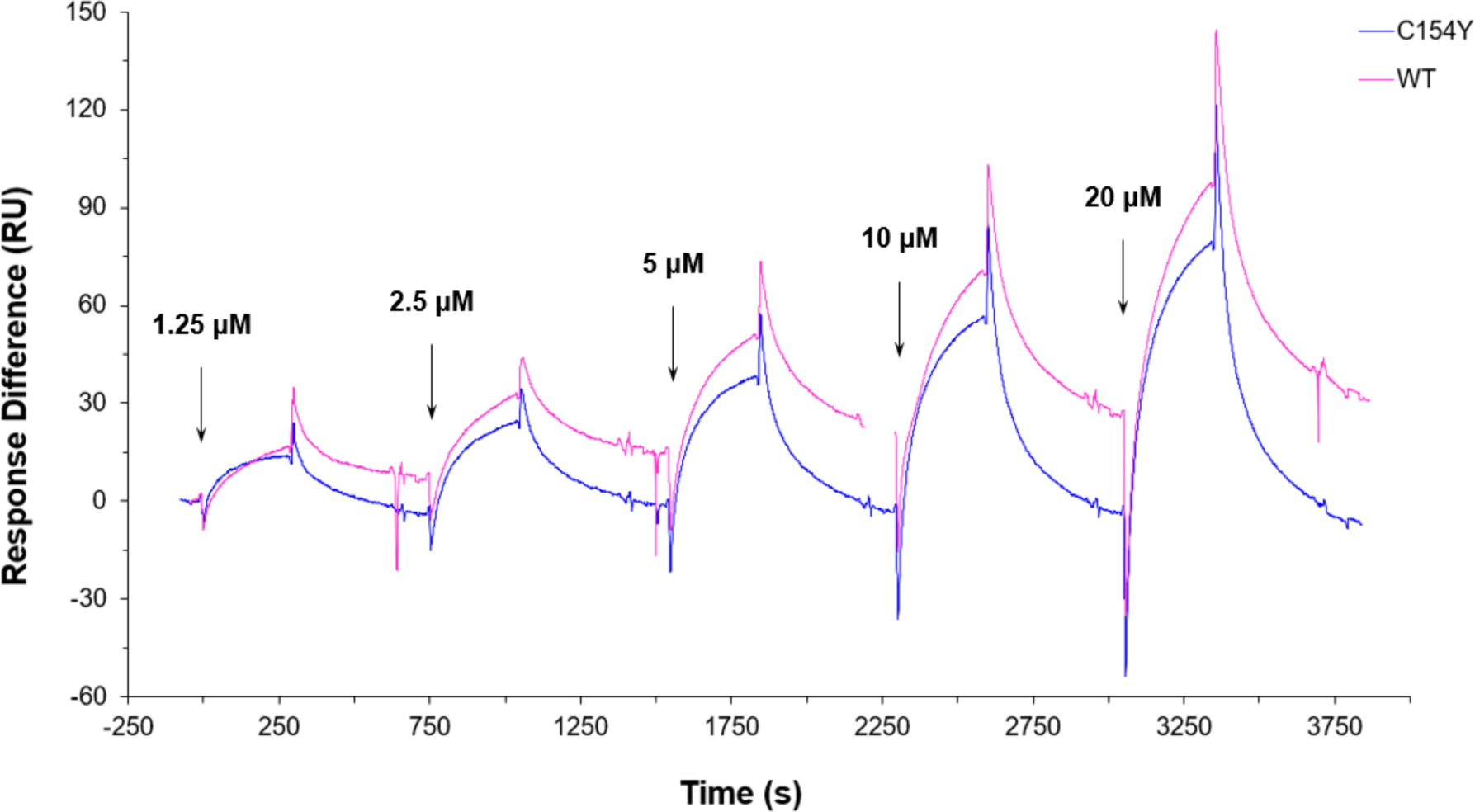
Surface plasmon resonance (SPR) sensorgram of the Kir2.1 WT (red) and Kir2.1-C154Y mutant (blue) (5844 RU level) immobilized onto a CM5 chip. Five sequential injections of PIP_2_ are shown, at 1.25-20 µM, and a flow of 5 µL·min^−1^. The sensorgram is expressed in response units (RU) relative to the time in seconds.

### Cryo-EM structure of the CTD domain of the Kir2.1-R312H mutant

The structure of the Kir2.1-R312H mutant was resolved by single-particle cryo-EM analysis to an average resolution of 6.0Å (Figure 6). Despite the medium resolution cryo-EM map, it is possible to observe that the model contains the TMD and CTD, and preserves all the typical structural features of Kir2.1 channels. Previously described (12) in a tetrameric arrangement. Therefore, R312H mutation does not affect the association of the subunits to form the correct tetrameric structure.

**Figure 6.**
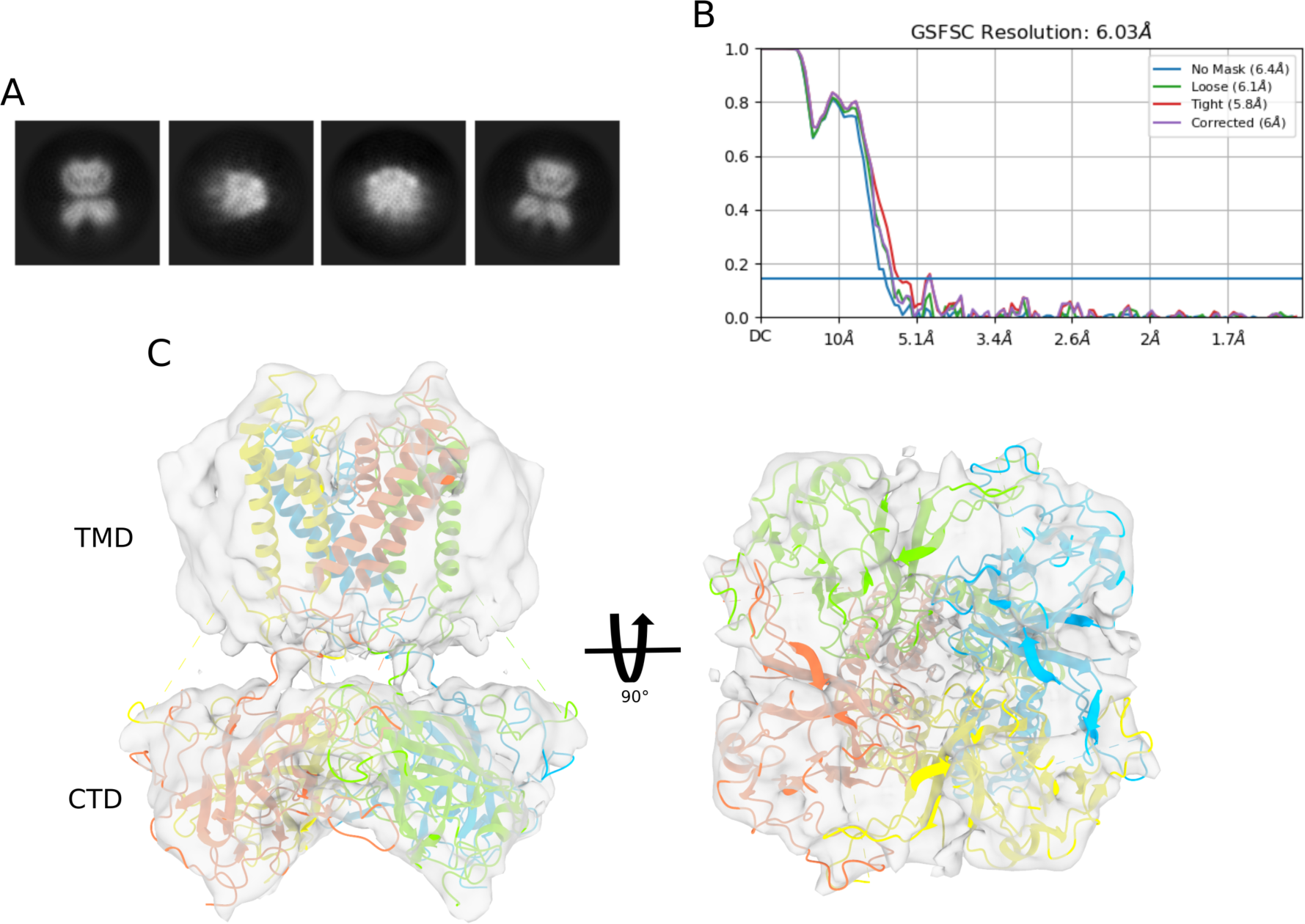
Cryo-EM structure of Kir2.1-R312H mutant A. Representative 2D-class averages. **B.** Fourier-shell correlation (FSC) curves of the final map reconstruction provided by CryoSPARC v4.4 **C.** Side and bottom views of the atomic structure of the Kir2.1-R312H mutant fitted in the cryo-map. The four chains (A to D) are highlighted as orange, red, blue and yellow cartoon respectively. TMD: transmembrane domain; CTD: cytoplasmic domain.

### MD simulations on Kir2.1-WT and Kir2.1-C154Y mutants

Structural analysis of the C122/C154 disulfide bond reveals its role in establishing a structural connection from the selectivity filter and the 147-153 loop to the extracellular loops (Figure 7A), which may be disrupted when the disulfide bond is broken (Figure 7B). To evaluate the impact of the C154Y mutation on the structure of the Kir2.1 channel, we performed molecular dynamics (MD) simulations on the cryo-EM structure of Kir2.1- WT (PDB ID 7ZDZ and incorporating the C154Y mutation in all four subunits of the Kir2.1 tetramer (Chains ABCD_C154Y_). Moreover, we also explored hybrid tetrameric WT-C154Y structures by placing the C154Y mutation in i) three subunits (Chains ABC_C154Y_, Chain D_WT_), ii) two adjacent subunits (Chains AB_C154Y_, Chains CD_WT_), iii) two diagonally opposite subunits (Chains AC_C154Y_, Chains BD_WT_), and iv) one subunit (Chain A_C154Y_, Chains BCD_WT_). The protein structure was embedded in a 1-palmitoyl-2-oleoyl-sn-glycero-3-phosphocholine (POPC) lipid bilayer and the MD simulations were performed for 200 ns in triplicate for all constructions.

**Figure 7.**
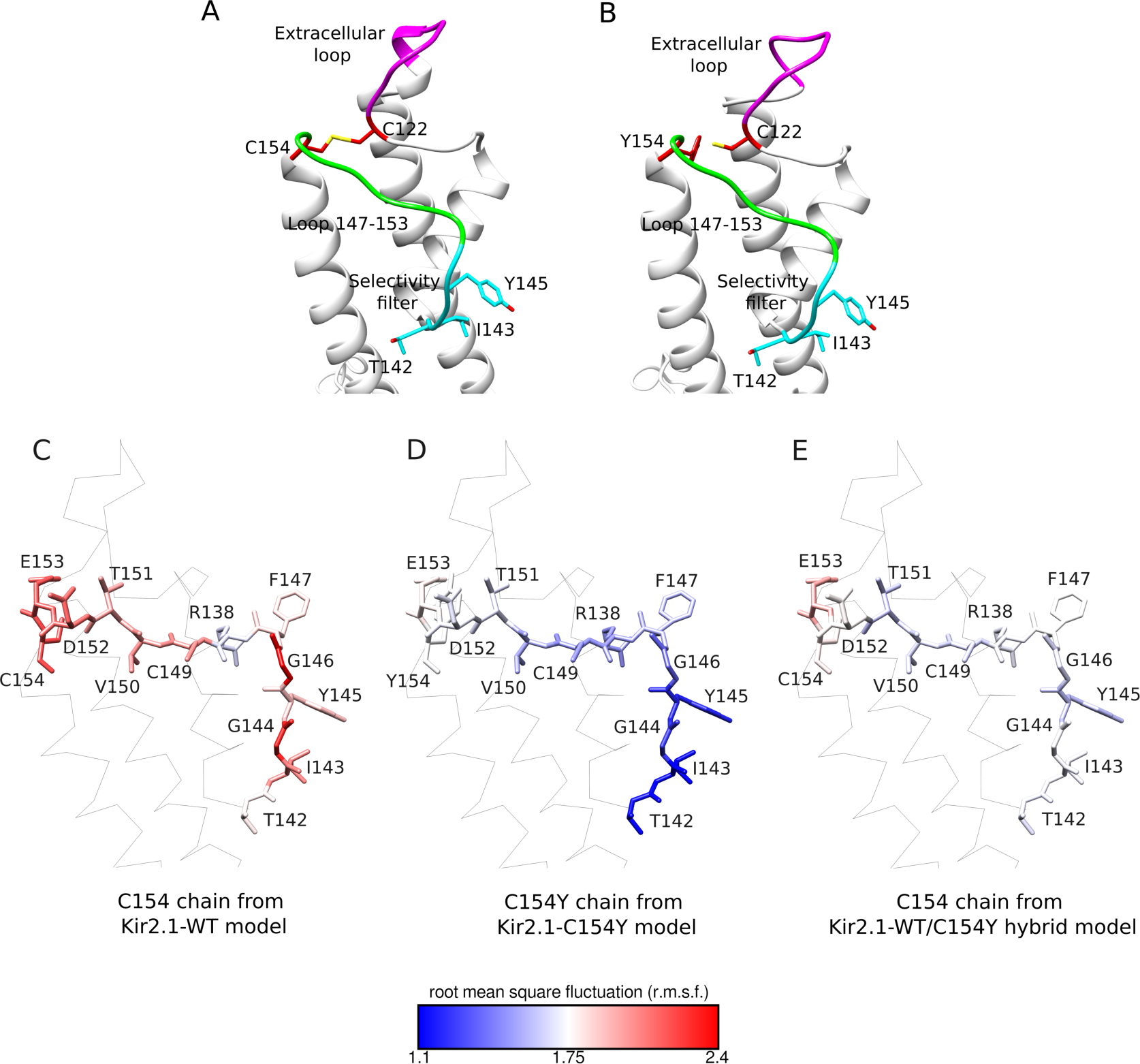
Analysis of the impact caused by the mutation C154Y on the Kir2.1 structure and on the root mean square fluctuation (rmsf) of the selectivity filter (region 142-146), the 147-153 loop, and the residue located at position 154. A. Kir2.1WT **B.** Kir2.1-C154Y Cartoon representation of the C122/C154 disulfide bond or C154Y and C122 are in red sticks. The extracellular loop (magenta) and the selectivity filter (cyan) are connected through the 146- 153 loop (green). The residues T142, I143, and Y145 from the selectivity filter are highlighted in cyan sticks. **C.** A representative WT chain (C154) from a representative replica of MD simulations performed with the Kir2.1-WT cryo-EM structure (PDB ID 7ZDZ). **D.** A representative C154Y chain from a representative replica of MD simulations performed after incorporating the C154Y in the four subunits of the Kir2.1 tetramer. **E.** A representative WT chain (C154) from a representative replica of MD simulations performed with structures containing the assembly of both WT and C154Y mutated subunits in the Kir2.1 tetramer (hybrid WT-C154Y models). In panels C, D, and E, the backbone a segment of the Kir2.1 structure containing part of the transmembrane domain (TMD) and the extracellular loop is shown in gray. The residues on which the rmsf was evaluated are showed in sticks and labeled according to their names and position. These residues are colored based on their rmsf values which are represented on the color scale ranging from 1.1 (blue, less flexible) to 2.4 (red, more flexible). Although a representative chain from a representative replica is shown in panels C, D and E, these results were consistent across all chains from all replicas performed.

Firstly, we assessed the impact of the C154Y mutation on the root mean square fluctuation (rmsf) of residue 154, the 147-153 loop, and the selectivity filter (residues 142-146) along the MD simulations (Figure 7C-E and Table 1). The all-WT tetramers showed rmsf values of 2.22 ± 0.14 at C154, 2.01 ± 0.12 at the 147-153 loop, and 2.02 ± 0.18 at the selectivity filter. The all-C154Y structures showed a decrease in rmsf for Y154 (1.74 ± 0.17), 147-153 loop (1.63 ± 0.13), and selectivity filter (1.23 ± 0.05), with a more pronounced effect on the latter one (Figure 7C-E and Table 1). The extracellular loops are the most flexible regions of the Kir2.1 structure, as demonstrated by our previous MD simulations results (12). The decrease in rmsf of the 147-163 loop and the selectivity filter in the C154Y structure suggests a decrease in the flexibility of the selectivity filter when the C122/C154 bond is disrupted, suggesting that the break of the C122/C154 disulfide bond interrupts the structural connection between the extracellular loop and the selectivity filter. Interestingly, in the MD simulations performed with the hybrid WT-C154Y structures, the WT subunits also exhibit a decrease of the rmsf of the 147-153 region (∼1.7-1.9) and selectivity filter (1.6-1.8) (Figure 7C-E and Table 1). These data imply that the impact of mutation in one subunit can affect the others, highlighting the required cooperativity of the selectivity filter in the tetramer to allow the flow of K^+^ ions.

**Table 1.**
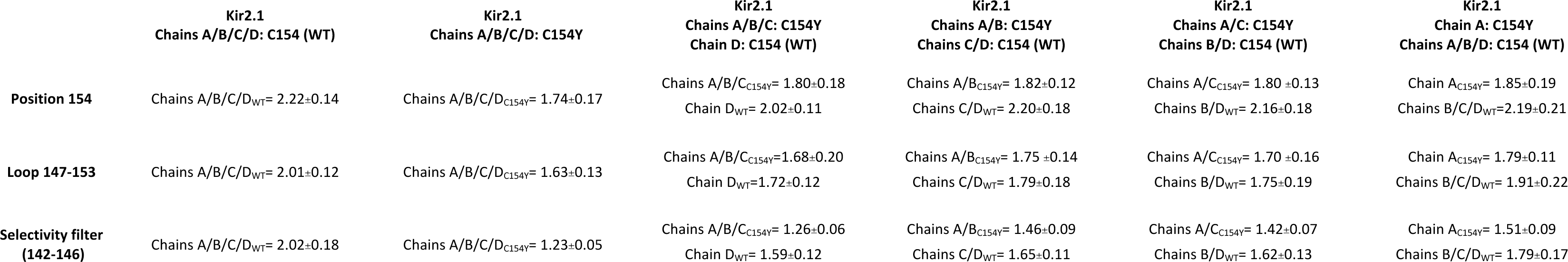
Average values of root mean square fluctuation (rmsf) calculated along the 200 ns of MD simulations in triplicate of the Kir2.1 channel cryo-EM structure WT (four chains C154) and after incorporating the C154Y mutation in the four subunits of the tetramer (Chains ABCD_C154Y_), in three subunits (Chains ABC_C154Y_, Chain D_WT_), in two adjacent (side-by-side) subunits (Chains AB_C154Y_, Chains CD_WT_), in two diagonally opposite subunits (Chains AC_C154Y_, Chains BD_WT_), and in one subunit (Chain A_C154Y_, Chains BCD_WT_).

Then, we investigated in detail the effect of breaking the disulfide bond on the selectivity filter structure. In a previous study on KcsA channels, it was demonstrated that the loss of flexibility of the selectivity filter leads to non-functional K^+^ channels. This loss prevents the attainment of dihedral angle values of the amino acids from this region that would favor the K^+^ ion flow (21). Then, we assessed the impact of the decreased rmsf values, caused by the C154Y mutation, by evaluating the occurrence frequency of dihedral angles of the selectivity filter residues: T142, I143, and Y145 along the MD simulations (Figure 8, Table 2, and Figures S6, S7 and S8). The MD simulations performed with Kir2.1-WT and Kir2.1-C154Y mutant revealed three main populations of dihedral angle frequencies for residues T142 and Y145. For residue T142, the most frequent dihedral angles range from −40° to −30° (brown), −20° to −10° (cyan), and 30° to 40° (green) (Figure 8A); whereas for Y145, dihedral angles range from −30° to −20° (brown), −10° to 0° (cyan), and 30° to 40° (green) (Figure 8B). In the MD simulations performed with the Kir2.1-C154Y mutant, a significant decrease in the frequency of the dihedral angles in the 30° to 40° range is observed in both residues (Figure 8A-B, Table 2, and Figures S6 and S7).

**Figure 8.**
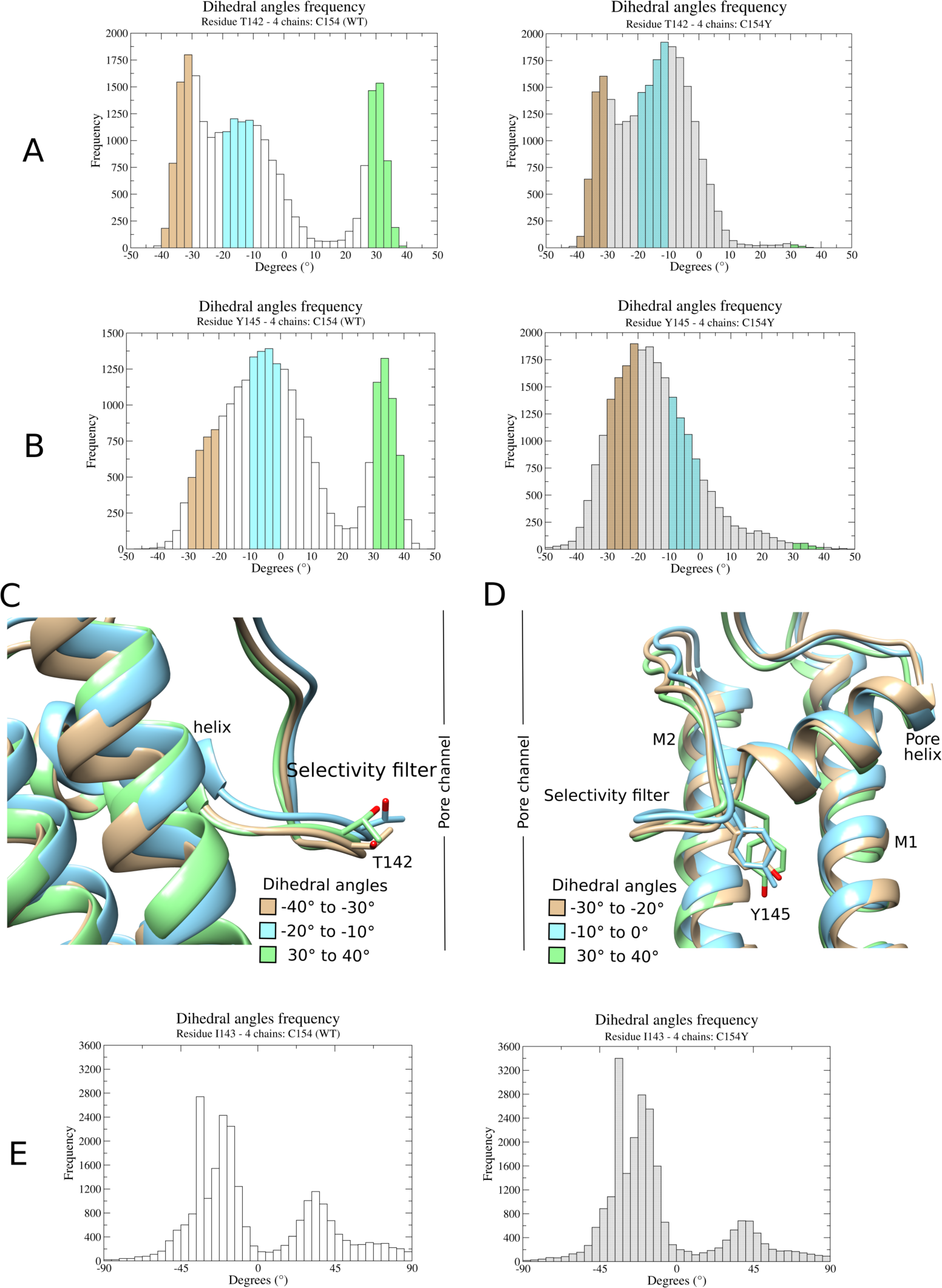
Dihedral angles analysis of the selectivity filter residues (T142, I143, and Y145) of the Kir2.1 channel. Dihedral angle frequency for the T142 (**panel A**), Y145 (**panel B**), and I143 (**panel E**) residues obtained along the 200 ns of MD simulations in triplicate of the Kir2.1 channel cryo-EM structure WT (four chains C154; left column) and after incorporating the C154Y mutation in the four subunits of the tetramer (Chains ABCD_C154Y_; right column). For residue T142 (**panel A**), the frequency of dihedral angles in the ranges of −40° to −30°, −20° to −10°, and 30° to 40° are highlighted in brown, cyan, and green, respectively. For residue Y145 (**panel B**), the frequency of dihedral angles in the ranges of −30° to −20°, −10° to 0°, and 30° to 40° are highlighted in brown, cyan, and green, respectively. **Panels C and D** show snapshots captured along the MD simulations to illustrate these dihedral angle ranges for the side chains (shown in sticks) of the residues T142 (**panel C**) and Y145 (**panel D**). The dihedral angle frequency analysis of residue I143 (**panel E**) is not highlighted because the impact of the C154Y mutation on it was inconclusive.

**Table 2.**
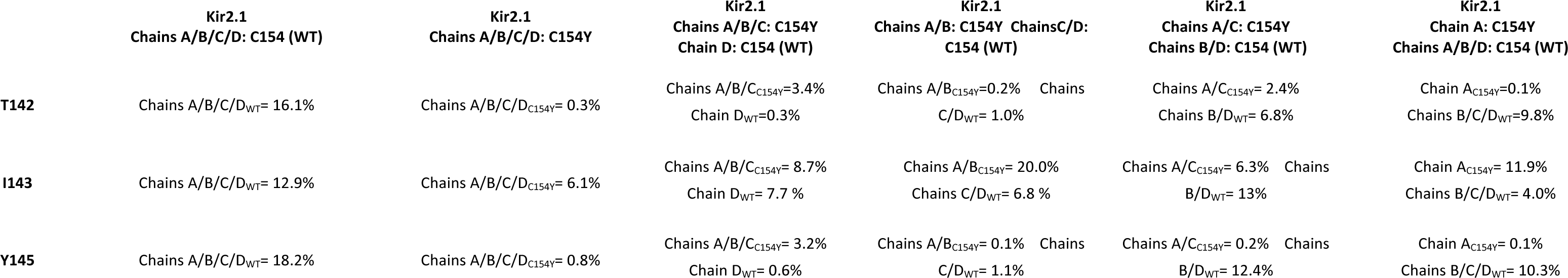
Frequency of dihedral angles in the range of 30°—40° for the T142, I143, and Y145 residues. These values were obtained along the 200 ns of MD simulations in triplicate of the Kir2.1 channel cryo-EM structure WT (four chains C154) and after incorporating the C154Y mutation in the four subunits of the tetramer (Chains ABCD_C154Y_), in three subunits (Chains ABC_C154Y_, Chain D_WT_), in two adjacent (side-by-side) subunits (Chains AB_C154Y_, Chains CD_WT_), in two diagonally opposite subunits (Chains AC_C154Y_, Chains BD_WT_), and in one subunit (Chain A_C154Y_, Chains BCD_WT_).

|  | Kir2.1<br>Chains A/B/C/D: C154 (WT) | Kir2.1<br>Chains A/B/C/D: C154Y | Kir2.1<br>Chains A/B/C: C154Y<br>Chain D: C154 (WT) | Kir2.1<br>Chains A/B: C154Y ChainsC/D:<br>C154 (WT) | Kir2.1<br>Chains A/C: C154Y<br>Chains B/D: C154 (WT) | Kir2.1<br>Chain A: C154Y<br>Chains A/B/D: C154 (WT) |
| --- | --- | --- | --- | --- | --- | --- |
| **T142** | Chains A/B/C/D<sub>WT</sub>= 16.1% | Chains A/B/C/D<sub>C154Y</sub>= 0.3% | Chains A/B/C<sub>C154Y</sub>=3.4%<br>Chain D<sub>WT</sub>=0.3% | Chains A/B<sub>C154Y</sub>=0.2%<br>Chains C/D<sub>WT</sub>= 1.0% | Chains A/C<sub>C154Y</sub>= 2.4%<br>Chains B/D<sub>WT</sub>= 6.8% | Chain A<sub>C154Y</sub>=0.1%<br>Chains B/C/D<sub>WT</sub>=9.8% |
| **I143** | Chains A/B/C/D<sub>WT</sub>= 12.9% | Chains A/B/C/D<sub>C154Y</sub>= 6.1% | Chains A/B/C<sub>C154Y</sub>= 8.7%<br>Chain D<sub>WT</sub>= 7.7 % | Chains A/B<sub>C154Y</sub>= 20.0%<br>Chains C/D<sub>WT</sub>= 6.8 % | Chains A/C<sub>C154Y</sub>= 6.3%<br>Chains B/D<sub>WT</sub>= 13% | Chain A<sub>C154Y</sub>= 11.9%<br>Chains B/C/D<sub>WT</sub>= 4.0% |
| **Y145** | Chains A/B/C/D<sub>WT</sub>= 18.2% | Chains A/B/C/D<sub>C154Y</sub>= 0.8% | Chains A/B/C<sub>C154Y</sub>= 3.2%<br>Chain D<sub>WT</sub>= 0.6% | Chains A/B<sub>C154Y</sub>= 0.1%<br>Chains C/D<sub>WT</sub>= 1.1% | Chains A/C<sub>C154Y</sub>= 0.2%<br>Chains B/D<sub>WT</sub>= 12.4% | Chain A<sub>C154Y</sub>= 0.1%<br>Chains B/C/D<sub>WT</sub>= 10.3% |

The WT subunits from the all-WT structure exhibit 16% and 18% frequency of a dihedral angle within the 30° to 40° range for the T142 and Y145 residues, respectively. In contrast, the C154Y-containing subunits show a decreased frequency, ranging from 0.1 to 3.4% (Table 2). This reduction in the frequency of dihedral angles within the 30° to 40° range for these residues in the chains containing the C154Y mutation is counterbalanced by an increase in the frequency of dihedral angles within the −20° to −10° range. Interestingly, this decrease in dihedral angle frequency in the 30° to 40° range is also observed for the T142 and Y145 residues of the WT subunits in the hybrid WT-C154Y structures. However, this effect is less pronounced in the hybrid structures where the C154Y mutation was placed in two diagonally opposite subunits (Chains AC_C154Y_, Chains BD_WT_), and in one subunit (Chain A_C154Y_, Chains BCD_WT_) (Table 2 and Figures S6 and S7). The orientation of the side chains of the T142 and Y145 residues in the three main populations of dihedral angle frequencies observed along MD simulations is illustrated in Figure 8C-D.

Concerning I143, in the Kir2.1-WT structure, and the Kir2.1-C154Y mutant, the most frequent dihedral angles range from −45° to −30° and 30° to 40° (Figure 8E). However, the mutant structure (all four C154Y subunits) shows a decrease in the frequency of dihedral angles within the 30° to 40° range compared to the WT subunits. Note that the impact in the dihedral angle frequencies is less pronounced than in the T142 and Y145 residues (Table 2 and Figure 8E). However, this phenomenon is not evident when analyzing the hybrid WT-C154Y structures (Table 2 and Figure S8). Therefore, the impact of the C154Y mutation on the dihedral angles of the I143 residue remains inconclusive.

Additionally, given the potential steric hindrance caused by the C154Y mutation, we explored the effect of the C154S mutation placed on four chains of the Kir2.1 structure on the plasticity of the selectivity filter and frequency of dihedral angles after 200 ns MD simulations in triplicate. The results for C154S were similar to those obtained for C154Y (Figure S9A and Table S1). Previous attempts to restore interactions between C122 and C154 by replacing these residues with aspartate and lysine, respectively, did not rescue the channel function (16). Here, we attempted to restore these interactions by replacing C122 and C154 with tyrosines (C122Y/C154Y) or tryptophans (C122W/C154W), by exploring the potential of the side chains in establishing aromatic-aromatic interactions after 200 ns MD simulations in triplicate. We did not observe any rescue of the rmsf or the dihedral angle frequency in the 30° to 40° range for the T142 and Y145 residues (Figure S9B-C and Table S1). These data indicate that only strong interactions between C122 and C154 residues, such as a covalent bond established in a disulfide bridge, would be able to maintain the structural connection between the extracellular loops and the selectivity filter.

## Discussion

The Kir2.1 channel undergoes many steps before it can conduct K^+^ ions. It must be properly recognized and targeted to the endoplasmic reticulum, folded, tetramerized, and trafficked to the plasma membrane. The mature channel must be able to interact with its modulators and effectively gate: open and close. The open state is triggered by the binding of PIP_2_ that induces several conformational changes in the channel (3, 18, 22). Any genetic alteration in the DNA coding region can hamper any of these stages, affect the function and dynamics of the protein, and lead to pathology. A more comprehensive understanding of the molecular mechanisms leading to dysfunction has the potential to improve the diagnosis and treatment of human genetic disorders.

### The dominant negative effect of the C154Y mutation arising from loss in the structural plasticity of the selectivity buffer

C154, a residue conserved in all eukaryotic Kir members, is located in the N-terminal of the M2 helix at the membrane interface. This residue establishes a disulfide bond with C122, which connects the extracellular loops to the selectivity filter towards the 147-153 loop (Figure 1B-C ). Three mutations at the C154 residue are responsible for ATS in patients, with a loss of function of the channel: C154F (23) C154Y (14), and C154G (ClinVar RCV001901246.2).

Our studies show that expression of the mutant C154Y in *X. laevis* oocytes resulted in no detectable currents (Figure 2). This support previous studies evaluating the C154A and C154S mutations which showed that these mutations abolished the current in *Xenopus laevis* oocytes (15, 16). Our results did not show that the C154Y mutation impacted the trafficking to the cell membrane, as shown with our immuno-fluorescence experiments in *X. laevis* oocytes and analysis of surface biotinylation of membrane proteins on *P. pastoris* cells (Figure 3A-B). Upon expression and purification, the Kir2.1-C154Y mutant exhibited reduced structural stability and increased susceptibility for aggregation (Figure S2). However, binding to PIP_2_ is still observed (Figure 5). The protein was not purified in sufficient amount to perform SEC-Malls and cryo-EM experiments. The aggregation phenomenon observed *in vitro* may not be present *in vivo*. When Kir2.1 is embedded in a cell membrane, only C122, C149, and C154 residues, located in the extracellular region are capable of forming disulfide bonds. Upon solubilization in DDM, the previously inaccessible intracellular cysteines become theoretically accessible to form disulfide bonds with the free C122 in the vicinity, potentially leading to protein aggregation *in vitro*. Moreover, the crowed cell environment, may provide different types of interactions (24) that can contribute to the stability of this mutant *in vivo*.

For the wild-type Kir2.1 channel, attempts to disrupt this disulfide bond by applying reducing agents (10-20 mM DTT or 10 mM reduced glutathione) extracellularly to cells expressing Kir2.1 showed minimal impact on channel currents (15, 16). However, the crystal structure of the Kir2.2 channel was obtained in the presence of 20 mM DTT and 3 mM TCEP reducing agents, and this disulfide bond remained unaffected (18). This suggested that the C122/C154 disulfide bond can withstand moderate concentrations of reducing agents and that this disulfide bond should be significant for the channel function (18). Mutations in residues C122 and C154 to serine in Kir2.3 channels (C113 and C145 according to Kir2.3 numbering) provided the same results in *X. laevis* oocytes: the mutant channels were expressed, assumed to be processed into tetramers, and trafficked to the membrane, but in a non-functional form (19). Whole-cell and single-channel currents in *X. laevis* oocytes of C122S and C154S mutants in Kir2.1 channels revealed that the presence of a single mutant subunit was enough to abolish the function of the tetrameric channel (16). Similarly, studies of the Kir2.1-C154F mutant in HEK293 cells demonstrated that this mutant traffics to the membrane and also exerts a dominant negative effect (23).

To understand the molecular basis for this possible dominant negative effect, we explored the structural properties of the region around the C154 in the Kir2.1 structure. Our MD simulation indicate that the disruption of the C122/C154 disulfide bond interrupts the structural connection between the flexible extracellular loops and the selectivity filter, resulting in a loss of structural plasticity of the selectivity filter and impairing the K^+^ flow. Indeed, C154Y mutation resulted in a decrease in the rmsf of the selectivity filter and a reduction in the frequency of dihedral angles in the 30° to 40° range for the T142 and Y145 residues located in this region (Tables 1 and 2; Figures 8, 9, S7 and S8). Interestingly, mutation of C154 to the flexible residues alanine or serine did not recover the current in Kir channels (15, 16). Our MD simulations of the C154S mutation placed in four chains of the Kir2.1 structure provided similar results in terms of the loss of structural plasticity of the selectivity filter (Figure S9A and Table S1). This indicates that only the flexibility level provided by the extracellular loops can provide the proper structural plasticity of the selectivity filter to allow the K^+^ flow.

Notably, the decrease of rmsf of the selectivity filter and in dihedral angles in the 30° to 40° range for the T142 and Y145 residues was observed even in the WT subunits of the hybrid WT-C154Y structures (Tables 1 and 2; Figures 8, 9, S6 and S7). These data highlight the cooperative role of the four subunits in coordinating the K^+^ flow at the selectivity filter. Thus, only one single mutated subunit can affect the tetrameric complex. In other words, for the C154Y mutation, all the subunits forming the channel must be functional to form an ion-conducting pore. Similar behavior was observed in HCN1 protein with a mutation at the pore (25). The presence of one such dominant-negative monomeric subunit suffices to render the entire channel complex nonfunctional. This a characteristic feature of mutations with dominant-negative effect (26, 27). Based on a probability model previously proposed for mutations with dominant-negative effect in homomeric complexes (26), we proposed a model of the association between C154Y and WT subunits in hybrids WT-C154Y Kir2.1 complexes (Figure 9A). According to this model, a unique mutated subunit disrupts the function of the entire complex and can lead to a loss of function of 93.75%. Note that this value is correct if the WT and mutated subunits are expressed in the same amount. In our case, when Kir2.1-C154Y is co-expressed with Kir2.1-WT we observed a loss of function of 70% (100% being the current registered for injection of 50 ng of Kir2.1-WT) (Figure 2B). The slight discrepancy between the two values coud be explained by the fact that, although the same amount of Kir2.1-WT and Kir2.1-C154Y cDNA were injected, Kir2.1-WT could be more readily expressed than Kir2.1- C154Y. With this assumption, more WT subunits would be available for association, leading to a higher current than expected.

**Figure 9.**
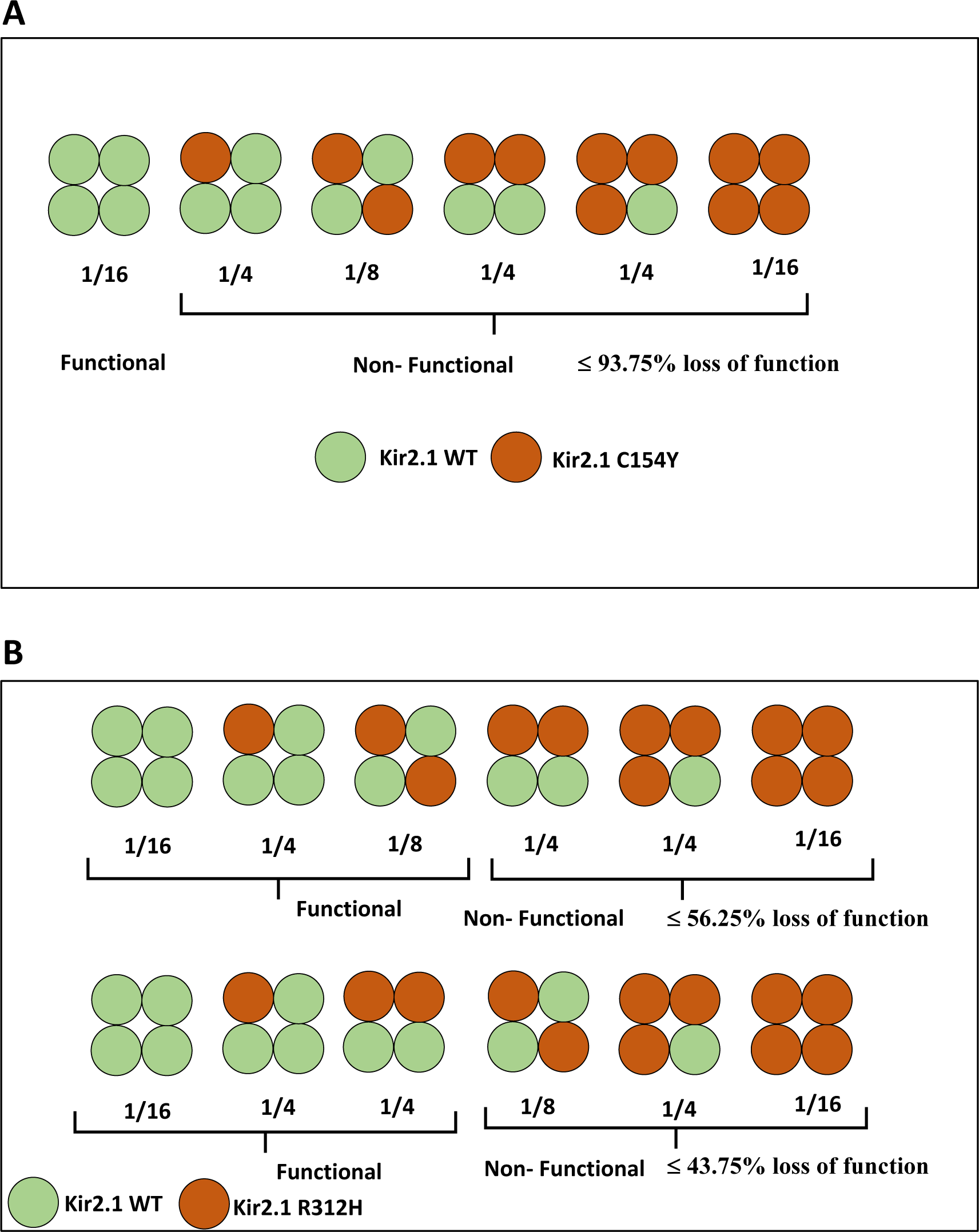
**Probabilistic models of the effect of association between WT and mutant subunits in Kir2.1 tetrameric complexes**. **A.** Model of the association between C154Y with WT subunits. This model predicts a dominant-negative effect of the C154Y mutation. One single C154Y mutant subunit blocks the activity of the entire complex, resulting in a loss-of-function of ≤ 93.75%. **B.** Model of the association between R312H and WT subunits. Two different propositions are presented in the upper and lower panels, respectively. The upper panel predicts a theoretical loss of function of ≤ 56.25% when R312H mutant subunits are placed side-by-side with WT subunits in the Kir2.1 tetramer. The lower panel predicts a theoretical loss of function of ≤ 43.75% when the R312H mutant subunits are placed diagonally opposed to WT subunits in the Kir2.1. These models were derived from a previously proposed probabilistic model (26)

### Effect of the mutation R312H on the gating mechanism

The residue R312 is located near the G-loop and the PIP2 binding site and mutant R312H is responsible for ATS disease (loss of function). Upon expression and purification, the Kir2.1- R312H mutant biochemical data (SEC and SEC-MALLS-RI) showed that it forms a homogeneous tetramer, indicating that this mutation did not impair the correct oligomerization of the channel (Figures 4 and S2). These data are also supported by our cryo-EM structure (Figure 6).

Immuno-fluorescence experiments in *X. laevis* oocytes and analysis of surface biotinylation of membrane proteins on *P. pastoris* cells show that R312H mutants are expressed and trafficked toward the cell membrane (Figure 3A-B). Expression of this mutant in *X. laevis* oocytes resulted in no detectable currents (Figure 2), although this mutation did not significantly impact the PIP_2_ binding as previously shown (12). The coexpression of Kir2.1-WT and R312H mutant in *X. laevis* oocytes resulted in a partial recovery of channel current, indicating that the dysfunction caused by R312H mutation can be partially rescued by association with the WT counterpart (Figure 2A).

Based on a probability model previously proposed (26), we modeled the effect of all combinations on the association of R312 and WT subunits along with their respective loss-of-function probabilities (Figure 9B). The difference in the loss-of-function probability arises from the position of the two mutated subunits within the tetramer complex. If side-by-side mutant subunits are non-functional, a theoretical loss of function of 56.25% is predicted (Figure 9B, upper panel). If diagonally opposed mutant subunits are dysfunctional, then the probability of loss of function is 43.75% (Figure 9B, lower panel). Alternatively, when any positioning of two mutant subunits results in non-functionality, the loss-of-function probability increases to 68.75%. Our electrophysiology results show a 40% loss of function when R312H and WT subunits are co-expressed compared to when only WT subunits are expressed (100% being the current registered for injection of 50 ng of Kir2.1-WT) (Figure 2A). Therefore, based on the loss-of-function prediction probabilities (Figure 9B), these results suggest that diagonally opposed mutant subunits lead to a loss of function, while side-by-side mutant subunits remain functional.

Our previous structural studies on Kir2.1 (12) describe a well-connected interaction network between the PIP_2_-binding site residues, R218 and K219, and the G-loop region (E303) via residues R312 and H221. Our previous data suggest that the conformational changes required for the G-loop opening are most likely controlled by PIP_2_ binding. Our previous *in silico* data show that the replacement of R312 with histidine leads to a complete loss of the interaction network described above, highlighting the fact that the structural integrity of this inter-subunit network of interactions, especially the E303-mediated salt bridges with R312 and H221 residues, is essential for the proper allosteric transmission of the signal between R312 and the G-loop of the adjacent subunit upon PIP_2_ binding and proper G-loop gating.

Based on partial recovery of current when Kir2.1-WT and R312H are coexpressed in *X. laevis* oocytes 1:1, our probability model, and our previous *in silico* results, we speculate that a partially open state of the channel at the G-loop could provide K^+^ current. We propose a structural hypothesis to understand this phenomenon (Figure S10). When the R312H mutation is placed in two diagonally opposite subunits, the G-loops of the R312H-containing subunits close the center of the channel pore, resulting in a nonfunctional channel. In contrast, when the R312H mutation is placed on side-by-side subunits, part of the channel pore remains unblocked, allowing the K^+^ flow even in this partially open state of the G-loop (Figure S10).

## Conclusion remarks

ATS poses a significant challenge in terms of therapeutic intervention as, to date, there is no effective treatment for this disease. Understanding molecular mechanisms can provide a pathway toward treatment. Therefore, understanding how ATS-causing mutations affect the Kir2.1 channel function at the molecular level provides a crucial foundation for designing targeted drugs to rescue the impaired function of these channels. Here, we were able to identify the molecular mechanisms by which two ATS-causing mutations (C154Y and R312H) affect the function of these channels. Both mutations did not affect the channel trafficking to the cell membrane; however, they impair the channel function, even though they can bind to the lipid activator PIP_2._ Notably, they hinder the channel function by different mechanisms. We also observed that C154Y has a more drastic effect on the Kir2.1 channels than the R312H loss-of-function mutation. Our data support that C154Y exerts a negative dominant effect, as the presence of this mutation in a single subunit of the tetramer is sufficient to render the entire complex nonfunctional. On the other hand, Kir2.1 channels containing R312H mutation in one or two adjacent subunits can still maintain channel current. The R312H mutation impairs the gating mechanism at the G-loop level, while the C154Y mutation impacts the K^+^ flow at the selectivity filter.

Our data can offer valuable insights that can guide potential strategies to rescue the function of both mutations. For the R312H mutation, gene therapy to increase the available wild-type protein would also result in increased currents and compounds targeting the enhancement of the stability in the intersubunit interaction network between E303, H221, and H312 could potentially correct the gating abnormalities and restore normal channel function. In the case of the dominant negative C154Y mutation, drug design should focus on the extracellular regions of the protein or directly on its selectivity filter. Compounds capable of reestablishing the structural connection between the extracellular loops and the selectivity filter, or providing increased flexibility and structural plasticity to the selectivity filter could potentially alleviate the hindrance to K^+^ flow caused by this mutation.

## Materials and methods

### hKir2.1 synthetic genes and expression in *Pichia pastoris* cells and *Xenopus laevis* oocytes

#### Protein expression in *Pichia pastoris*

Protein expression was handled as previously described (12, 28). Briefly, synthetic KCNJ2 genes encoding residues 1 to 427 (the whole sequence Uniprot reference P63252) of human Kir2.1 WT, R312H, or C154Y mutants optimized for expression in yeast were cloned in a pPIC9K vector upstream of a sequence coding for a PreScission protease cleavage site (LEVLFQGP) followed by a linker of 11 amino acids and a 10His tag (GenScript). The plasmids were introduced in *Pichia pastoris* strain SMD1163 (*his4, pep4, prb1*), and the resulting colonies were further analyzed *via* an *in situ* Yeastern blot immunoassay in order to identify the best-expressing clones. Protein production was performed as previously described (12). Cells were harvested by centrifugation, washed in PBS buffer (pH 7.4), and stored at −80 °C until use.

#### Protein expression in *Xenopus laevis* oocytes

The hKir2.1 cDNA gene coding the sequence of human Kir2.1 was cloned into the mammalian expression vector pMT3 containing an Ampicillin resistance gene (GenScript). Kir2.1 pathological mutants (C154Y, R312H) were generated by polymerase chain reaction (PCR) with synthetic primers using the CloneAmp HiFi PCR Premix (Takara). Primer sets used for site-directed mutagenesis were the forward sequence ‘5- GTGACTGATGAG<u>TAC</u>CCAATTGCAGTGTTTATGGTG -3’, and the reverse sequence ‘5- CCACCATAAACACTGCAATTGG<u>GTA</u>CTCATCAGTCAC -3’ for C154Y, the forward sequence ‘5- CTCAGTGT<u>CAC</u>TCTAGCTATCTGGC -3’ and the reverse sequence ‘5- GCCAGATAGCTAGA<u>GTG</u>ACACTGAG -3’ for R312H, respectively (underlined nucleic acids indicate the mutated site). Mutations and DNA constructs were confirmed by sequencing (Eurofins, Cologne, Germany). *Xenopus laevis* oocytes at stage VI were provided by the University of Paris Saclay (TEFOR Paris-Saclay CNRS UAR2020/INRAE UMS 1451) and kept in Barth’s solution (87.34 mM NaCl, 1 mM KCl, 0.66 mM CaNO_3_, 0.75 mM CaCl_2_, 0.82 mM MgSO_4_, 2.4 mM NaHCO_3_, 10 mM HEPES and pH adjusted at 7.6 with NaOH). Oocytes were dissociated from ovary segments by digestion in 1 mg·mL^−1^ collagenase type II (Gibco) dissolved in OR2 solution (85 mM NaCl, 1 mM MgCl_2_, 5 mM HEPES, pH adjusted to 7.6 with KOH) for 1-2 h at room temperature under gentle agitation. Defolliculated oocytes were rinsed extensively with Barth’s solution and thereafter stored at 4 °C in Barth’s solution until required. Single oocytes were injected with 50 ng·μL^−1^ of cDNA coding for the monomeric sequence of hKir2.1, together with a cDNA encoding eGFP at 25 ng·μL^−1^, into the oocyte nucleus by air injection. Injected oocytes were incubated at 18 °C in Barth’s solution for 2-3 days for heterologous expression. In the case of co-expression of Kir2.1 wild-type and pathological mutants (C154Y, R312H), equal amounts (25 ng·μL^−1^) of mutants and wild-type cDNA were injected.

#### Two-electrode voltage clamp (TEVC) recording in *Xenopus laevis* oocytes

Macroscopic potassium currents (whole-cell currents) were recorded 2-3 days after injection of hKir2.1 cDNA (Genscript) using a two-electrode voltage-clamp amplifier (OC- 725C, Warner Instruments) and Clampex 10.6 software (Molecular Devices) for data acquisition. Oocytes were perfused at room temperature with KD10 solution (10 mM KCl, 88 mM NaCl, 1.8 mM CaCl_2_, 1 mM MgCl_2_, and 5 mM HEPES, pH adjusted to 7.4 with NaOH. Voltage-sensing and current-recording glass microelectrodes were filled with 3 M KCl and pulled to be used at resistances between 0.2-2 MΩ. Currents were elicited by 500 ms pulses from −150 to +30 mV in 10 mV increments from a holding potential of −60 mV. Recorded currents were digitized at 500 Hz using a Digidata 1550A interface (Molecular Devices) and filtered at 100 Hz. No leak subtraction was performed during the experiments. Data analysis was performed with Clampfit 10.6 software (Molecular Devices). In all figures, statistical data are presented as mean ± S.E.M., where n represents the number of oocytes recorded. Co-injection of the pMT3 plasmid carrying the eGFP fluorescent protein was used to check that the injection of the cDNA was performed successfully. Fluorescent oocytes were the indication of a successful injection of the plasmid.

#### Immunofluorescence experiments in *Xenopus laevis* oocytes

For oocyte immunofluorescence experiments, GFP-positive oocytes were fixed in 4% paraformaldehyde (PFA) in phosphate-buffered saline (PBS) overnight at 4 °C. Fixed oocytes were washed two times with PBS for 5 min each and then treated with 0.1% Triton X-100 in PBS for 10 min for permeabilization. Non-specific staining was blocked using 10% horse serum in PBS for 30 min at room temperature, followed by two washes with 2% horse serum in PBS for 5 min each. For immunolabeling, oocytes were incubated with the Rabbit anti-Kir2.1 primary antibody (1:200, Alomone Labs) in 2% horse serum for 1 h 30 min at room temperature and washed two times with 2% horse serum for 5 min each. Oocytes were labeled with the Cy3- conjugated anti-Rabbit secondary antibody (1:300, Invitrogen) in 2% horse serum for 1 h at room temperature and washed with 2% horse serum for 5 min. After 2 h fixation in 4% PFA in PBS at 4 °C, oocytes were placed in 3% low-gelling temperature agarose overnight at 4 °C. 40 µm slices of each oocyte were made using a Leica VT1000 S vibratome (Leica BioSystems). Several slices per oocyte (7-9 slices per two oocytes, three separate experiments) were mounted on a microscope slide with the ProLong™ Diamond Antifade Mountant (Invitrogen) and analyzed using an epi-fluorescence microscope with constant exposure time for visualization of cell surface expression. GFP-injected oocytes were used as negative controls. Oocytes injected with the pMT3 plasmid coding for the human glycine receptor (GlyR) were used as positive controls, following the same protocol as for hKir2.1, but with a mouse anti-GlyR primary antibody (1:500, Synaptic Systems) and a Cy3-conjugated anti-mouse (1:300, Life Technologies) secondary antibody.

#### Cell surface protein biotinylation of *Pichia pastoris* cells

The biotinylation assays consisted in labeling cell surface proteins with a biotin reagent before lysing the cells and purifying the targeted proteins. Briefly, Kir 2.1-WT and mutants were expressed in yeast cells (see above). Yeast pellet (4 g) was resuspended in 10 mL PBS buffer (137 mM NaCl, 2.7 mM KCl, pH 8, 8 mM Na_2_HPO_4,_ and 2 mM KH_2_PO_4_) supplemented with 10 mM of Sulfo-NHS-LC-Biotin (Thermo Fisher) and incubated for 30 min with agitation at room temperature. The cells were then washed three times with the PBS buffer and subjected to cell lysis, protein extraction, and purification as previously described (12). Proteins were analyzed on 10% gels SDS-PAGE either directly stained with Coomassie blue or electroblotted. Western blot membranes were incubated overnight at 4 °C, using either Extravidin-HRP (Sigma) at 1/1000 dilution in 10 ml of PBS-T buffer (PBS 1x, Tween20 0.02%, Milk 5%), targeting biotin, or an anti-Histidine conjugated with peroxidase at 1/1000 dilution in 10 mL of PBS-T buffer targeting His-tag Kir-2.1 (WT and mutants). The signal was revealed using the SuperSignal West PICO Plus kit (Thermo Fisher) and detected using Imager 680 (Amersham) with a 10-second exposition period.

### Protein solubilization and purification in DDM

Purification of Kir2.1-WT and mutants (R312H and C154Y) was performed in DDM (n-Dodecyl-β-D-maltoside, Glycon), as previously described (12). Briefly, yeast cells were ruptured using either Constant System Cell Disrupter or FastPrep 24 (MP Biomedicals). The membrane proteins were solubilized by adding 29.3 mM DDM (1.5%). The purification process involved an affinity chromatography step using cobalt affinity resin (TALON, Clontech), followed by a size exclusion chromatography (SEC) on either a Superdex® 200 (10/300) GL column or Superdex® 200 Increase (10/300) column (Cytiva) pre-equilibrated with TKED buffer (20 mM Tris-HCl pH 7.4, 150 mM KCl, 1 mM EDTA, 0.03%/0.59 mM DDM) using the Äkta Purifier system (Cytiva). Fractions corresponding to the tetramer were pooled, 2 mM DTT, was added, and concentrated to 0.7–1 mg·mL^−1^. The solubilization in other detergents is detailed in the supplementary material. For cryo-EM, the protein was solubilized in DDM and exchanged with amphipol A8-35 before SEC as described previously (29).

### SEC-MALLS-RI, a triple detection method to characterize membrane proteins in solution

SEC-MALLS-RI experiments were carried out on a Shimadzu HPLC coupled to an Optilab T-rEX refractometer and a miniDawn TREOS Multi-Angle Laser Light Scattering (MALLS) detector (both from Wyatt Technology). A Superdex 200 Increase column (Cytiva), equivalent to the S200-GL column but supporting high pressure, was used. The system was equilibrated overnight with the SEC mobile phase (20 mM Tris-HCl, pH 7.4, 150 mM KCl, 1 mM EDTA, and 0.03% DDM) at 0.3 mL·min^−1^ flow rate. The RI purge was set in on-mode to equilibrate the reference and measurement chambers. The temperature of the refractometer cell was fixed at 25 °C. The next day, the UV lamp and LS laser were turned on. Once the UV, LS, and RI baselines were stable, the mobile phase RI was noted, the RI purge was switched off, and the RI baseline was set to zero. 20 µL of BSA at 5 mg·mL^−1^ were injected to calculate the inter-detector volumes. The signal-acquisition interval was set to 0.125 s. The molar mass of molecules (membrane protein detergent complexes and detergent micelles) passing through the SEC column could then be determined without calibration by measuring their static light scattering and knowing the respective extinction coefficients and refractive index increments of the protein and the detergent, by using the module Protein Conjugate from the Astra V software (Wyatt Technology). The methodology of mass calculation is thoroughly described in (30) and detailed in the supplementary material.

#### Determination of the mass of the Kir2.1-DDM complex

First, Kir2.1 samples were analyzed following isolation of the estimated tetramer peak post-SEC purification, concentrated about 100 times with a 100 kDa cutoff filter. 20 µL of Kir2.1 at 0.8 mg·mL^−1^ in 0.03% DDM were injected and eluted at 0.4 mL·min^−1^ flow rate at room temperature. The best results were obtained when the protein was injected into SEC-MALLS just after affinity chromatography.

The flow rate was reduced from 0.4 mL·min^−1^ to 0.3 mL·min^−1^ and the column was systematically washed with NaOH to remove background noise coming from impurities fixed on the column. The same protocol was used for all other detergent/protein Kir2.1 mixtures.

### Surface plasmon resonance

The interaction between Kir2.1 and the lipid PIP_2_ was characterized by SPR on a Biacore 3000 instrument (Cytiva) controlled by Biacore 3000 Control software v4.1. The experiments were performed following the protocol described in (12). All biosensor experiments were performed in triplicate at 25 °C using the running buffer 20 mM Tris-HCl (pH 7.5), 150 mM KCl, 0.05 mM EDTA, and 0.05% DDM). In all experiments, a flow cell was left blank to be used as a reference for the sensorgrams. Kir2.1 (WT/C154Y) was immobilized onto a carboxymethylated dextran (CM5) sensor chip. The activation of CM5 chips and immobilization of the protein steps were done using standard Biacore procedures. The binding and kinetic assays were performed using single-cycle kinetics. PIP2 was serially diluted in running buffer to working concentrations (0.15 to 20 μM). For each cycle, PIP_2_ was injected at increasing concentration with a flow of 5 μl/min over both the reference cell and the ligand cell. Before the first PIP2 injection, a running buffer was injected and used as a double reference. Each injection consisted of 300-s contact time with 300-s dissociation time. No regeneration step was done between injections, as these buffers were detrimental to Kir2.1. The data were analyzed using BIA evaluation software 4.1, and kinetic parameters were determined using general fit and the titration kinetics 1:1 binding with drift model. The K_D_ was determined directly from the fitted curves.

### Sample preparation and cryo-EM data collection of Kir2.1-R312H mutant

Three microliters of Kir2.1 R312H mutant form were placed on glow-discharged (25 s) holey carbon-coated grids (Quantifoil R1.2/1.3, Au 200 mesh), blotted for 3.0 s, and flash-frozen in liquid ethane using a Vitrobot Mark III (Thermo Fisher Scientific) operated at 4 °C and 100% humidity. The EM data collection statistics are available in Table S2. A total of 10,762 movies were collected on a Titan Krios G4i microscope at Institut Pasteur (Paris) operated at 300 kV equipped with a Falcon 4 direct electron detector and a Selectrix Image Filter (Thermo Fisher Scientific). The automation of the data collection was done with the software EPU. Movies were recorded in electron-counting mode using EER fractionation at 165,000x nominal magnification with an exposure time of 2.93 s, and a total dose of 40 electrons/Å. A defocus range of −1.2 to −2.4 μm was used and two images were acquired per foil hole. The pixel size was 0.73 Å/pixel.

### Cryo-EM data processing of Kir2.1-R312H mutant CTD domain

The movies were motion-corrected and dose-weighted using MotionCor2 (31), and contrast function parameters (CTF) were estimated using CTFFIND4 (32). The corrected micrographs were filtered using a maximum CTF-fit resolution of 8 Å and a maximum total full-frame motion distance of 20 pixels. The image processing was then performed using CryoSPARC v4.4 (33). A blob picking followed by 2D classification was performed to generate templates for automated template-picking. After template-picking, the particles were submitted to three rounds of 2D classification to remove false picks, ice contamination, and classes with unclear features. A total of 146,539 particles were subjected to 3D *ab initio* reconstruction using no symmetry, followed by a single round of non-uniform refinement using C4 symmetry that provided a final map at 6.03 Å resolution (gold-standard FSC= 0.143). This final map was subjected to B-factor sharpening using the Auto-sharpen tool available in the PHENIX software suite (34). Cryo-EM data collection information is summarized in Table S2.

### Model build and refinement

The Kir2.1-WT structure was placed in the final sharpened cryo-EM map using the Dock in Map tool available in PHENIX (34). Once the model was placed in the cryo-EM density map, the R312H murtatuon was introduced using Coot (35). Due to the absence of clear cryo-EM electron density, the 61-70 loop was not modeled. Iterative cycles of refinment using the phenix.real_space_refine in PHENIX and manual adjustments in Coot provided the final cryo-EM model for the Kir2.1-R312H murtant. Refinment statistics are summarized in Table S2

### Molecular dynamics simulations of Kir2.1 C154Y mutant

To evaluate the impact of C154Y mutation on the structure and dynamics of Kir2.1 channels, this mutation was modeled on the cryo-EM structure of the human Kir2.1 channel (PDB ID 7ZDZ) (12) using the CHARMM-GUI server (36), followed by MD simulations employing NAMD software (37) under the CHARMM36m force field (38). For that, the mutation was placed in one monomer (Chain A_C154Y_), in two adjacent (side-by-side) monomers (Chains AB_C154Y_), in two diagonally opposite monomers (Chains AC_C154Y_), in three monomers (Chains ABC_C154Y_) and four monomers (Chains ABCD_C154Y_) of the Kir2.1 tetramer. Moreover, we also performed MD simulation with the WT and the C154S, C122Y/C154Y and C122W/C154W mutations placed on the four chains of the Kir2.1 structure. PROKTA (39) was used to assign the protonation states of the protein at pH7.0 using PARSE forcefield.

All preparation steps were performed using the CHARMM-GUI server (36). All mutant forms designed were embedded in a 100% POPC lipid bilayer and it was used a KCl concentration of 0.15 M was used to calculate the number of neutralizing atoms. The system was heated and equilibrated in the standard equilibration protocol suggested by CHARMM-GUI developers, gradually decreasing protein and membrane atomic positional restraints for 2 ns. Last, the production run was performed without any positional restraints for 200 ns (2000 frames; 0.1 ns per frame) in triplicate for all the systems studied (WT and the different built mutant forms of the Kir2.1 channel), resulting in a total of 27 MD simulations.

The dihedral angles of the T142, I143 and Y145 residues were calculated for the full trajectories using the command QUICK in CHARMM v43b1 (40) following the atom sequence available in the protein structure file (psf) obtained with the CHARMM-GUI: v, Cβ, Cγ, and C for T142; Cα, Cβ, Cδ, and C for I143; and Cα, Cβ, Cζ, and C for Y145. The root mean square fluctuation (rmsf) of Cα atoms was calculated over 200 ns after frames alignment using the last equilibrated frame as reference.

## Accession numbers

PDB 8QQL and EMD-18595

## Supporting information

Supplementary Figures tables and results

## Acknowledgments

We thank C. Travert and Dr. F. Skouri-Panet for GEMME biochemistry & geobiology facility at IMPMC; R. Boucher for his help on MD simulation; Nanoimaging Core facility (C2RT) at the Institut Pasteur and the help of J.-M. Winter, E. Salazar, S. Tachon, and M. Vos.

