## Supplementary Figures tables and results for "Biochemical, biophysical, and structural investigations of two mutants (C154Y and R312H) of the human Kir2.1 channel involved in the Andersen-Tawil syndrome"

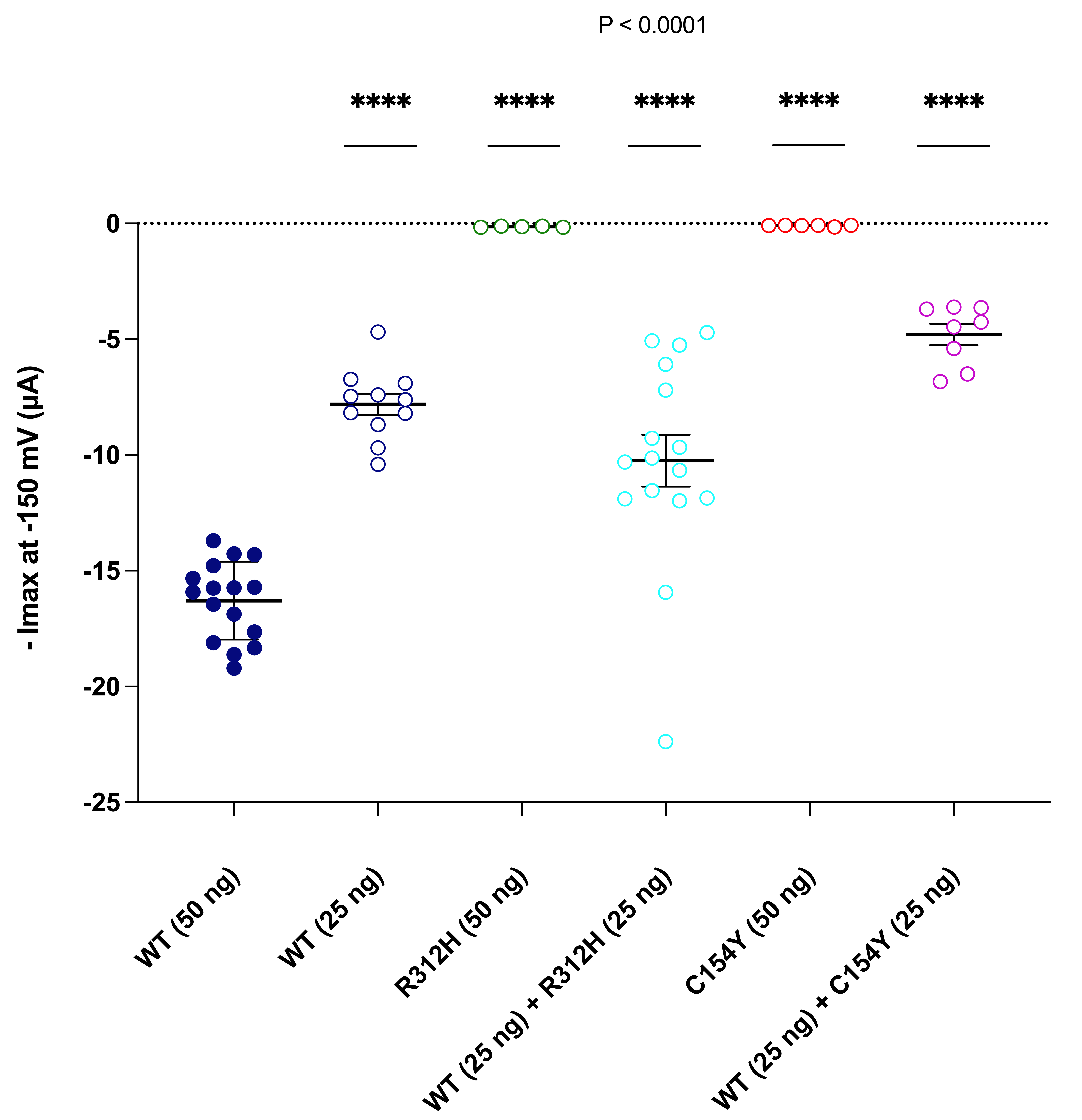

**Figure S1.** **Maximum negative current values at -150 mV of Kir2.1-WT, C154Y and R312H mutants recorded with TEVC in *Xenopus laevis* oocytes.** Unpaired two-sided student t-test indicates the significance of the effect of the mutations on -Imax at -150 mV (****P < 0.0001) (n=6-16 oocytes per group).

**A	B**

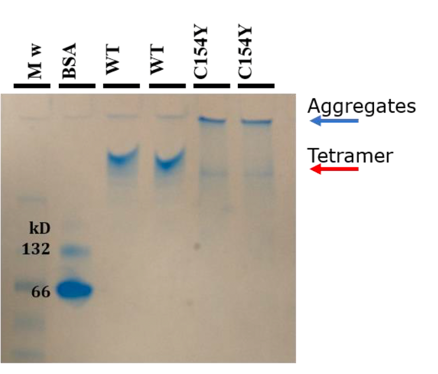

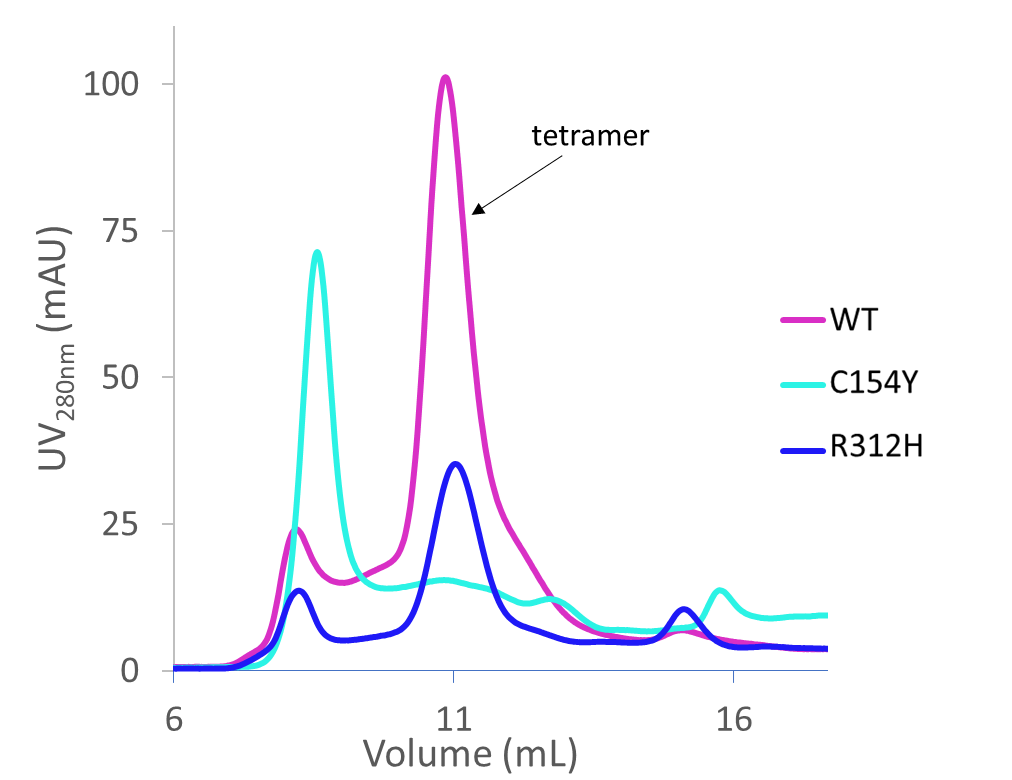

**Figure S2. Size-exclusion chromatography profile (SEC) of Kir2.1-WT and R312H and C154Y mutants. A.** Comparison between SEC profiles of Kir2.1-WT, and R312H and C154Y mutants on Superdex® 200 (10/300) GL column (Cytiva) **B.** Native PAGE gel comparing the migration of Kir2.1-WT and C154Y mutant. Kir2.1 WT migrates as single entity, corresponding to tetramers, positioned well above the BSA dimer. C154Y shows a band at the same level of tetramers and a more prominent one at the well level, corresponding to aggregates.

**SEC-MALL-RI:** **Kir2.1-WT solubilized in FC14 followed by an exchange in DDM or PCC-Malt detergents**

Materials and Methods

For extraction and purification using FC14 (tetradecylphosphocholine), the same protocol was used, but with a different buffer composition: 0.5% FC14 instead of 1.5% DDM for protein solubilization and 0.03% FC14 instead of 0.03% DDM for the purification steps. The exchange of Kir2.1 solubilized and purified in FC14 (Kir2.1-FC14) to DDM (Kir2.1-FC14-DDM) and PCC-Malt (4-trans-(4-trans-Propylcyclohexyl)-cyclohexyl maltoside from Glycon) (Kir2.1-FC14-PCCMalt) was performed as follow: after solubilization in 0.5% FC14, 20-30 CV of buffer containing DDM 0.05% or PCC-Malt 0.05% was added to the Kir2.1 immobilized in a cobalt affinity resin, followed by the usual elution and SEC steps in 0.03% DDM (Kir2.1 FC14-DDM) or 0.03% PCC-Malt(Kir2.1 FC14-PCC-Malt).

Results

Previous work demonstrated that functional FLAG-tagged Kir2.1 could be successfully isolated from S. cerevisiae cells using 1% FC14 (tetradecyl-phosphocholine) (1). For our construct, 0.5% FC14 provided excellent solubilization yields (Figure S3A). Purifying Kir2.1 exclusively in FC14 (0.03%) resulted in 55% of Kir2.1 being purified as tetramers (RH = 60 Å) and 43% eluting as monomers (RH = 31.3 Å) (Figure S3B, in yellow). Due to this high proportion of monomers, we attempted detergent exchanges with either 0.03% DDM (Kir2.1-FC14-DDM) or 0.03% PCC-Malt (4-trans-(4-trans-Propylcyclohexyl)-cyclohexyl α-maltoside) (Kir2.1-FC14-PCC-Malt). PCC-Malt is a detergent with a maltoside head like DDM, but the hydrophobic tail bears two cyclohexyls instead of a linear alkyl chain. It has been increasingly used recently and has been shown to increase membrane protein stability (2). After the detergent exchange, we recovered the SEC peak corresponding to the Kir2.1 tetramer (RH = 62.8 Å) (Figure S3B, in purple). However, this sample formed aggregates when concentrated by a factor of ~ 25 (Figure S3B, in red). We analyzed further the properties of Kir2.1-FC14-DDM samples using SEC-MALLS-RI. The profile is shown in Figure S4A and is similar to Kir2.1 WT in DDM (Figure 4A), with the tetrameric peak eluting at the expected volume (~9.7 mL). However, we observed two noticeable differences: i) although all three signals (LS, ∆RI, and UV) were visible, they did not align (Figure S4B) indicating that the sample does not appear homogeneous along the peak. Consequently, we could not determine the absolute protein molar mass with accuracy; and ii) a depletion in the detergent LS and ∆RI signals was observed at 15 mL suggesting that the DDM is pumped from the mobile phase to the protein complex and the extra micelles (Figure S4A, red and blue curves).

Protein-free FC14 micelles eluted at ~14 mL with an equal peak contribution from LS and ∆RI and minimal UV (Figure S4C). The depletion in the LS and ∆RI signals was also observed in the two controls, 5% FC14 and 5% FC14-DDM (Figure S4C). The detergent peak obtained in the sample (Figure S4A) eluted similarly to that of the FC14-DDM mixture, indicating an incomplete exchange of FC14 during the elution on the affinity column. Nonetheless, the depletion of DDM from the baselines implies that FC14 and DDM are miscible; therefore, a complete exchange could be possible under different exchange conditions, for instance, DDM concentrations should be significantly increased to displace FC14, which has a CMC slightly lower than DDM (0.12 mM and 0.17 mM respectively). Kir2.1-FC14-PCC-Malt samples were also investigated by SEC-MALLS-RI, and their profile was similar to that obtained with Kir2.1 FC14-DDM samples (Figure S5). The peak corresponding to Kir2.1 tetramers was obtained at the same elution volume (~9.7 mL) (Figure S5A). As for the Kir2.1-FC14-DDM samples, detergent signal depletion at 15 mL was also present (Figure S5A), although less pronounced than Kir2.1 FC14DDM. The three signals (LS, ∆RI and UV) did not align (Figure S5B) suggesting a detergent mixture around Kir2.1. The depletion in the LS and ∆RI signals was also observed in the two controls (5% FC14 and 5% FC14-PCC-Malt) (Figure S5C). These SEC-MALLS-RI experiments indicated an incomplete exchange of FC14 either with DDM or with PCC-Malt in Kir2.1 samples obtained from *Pichia pastoris* cells.

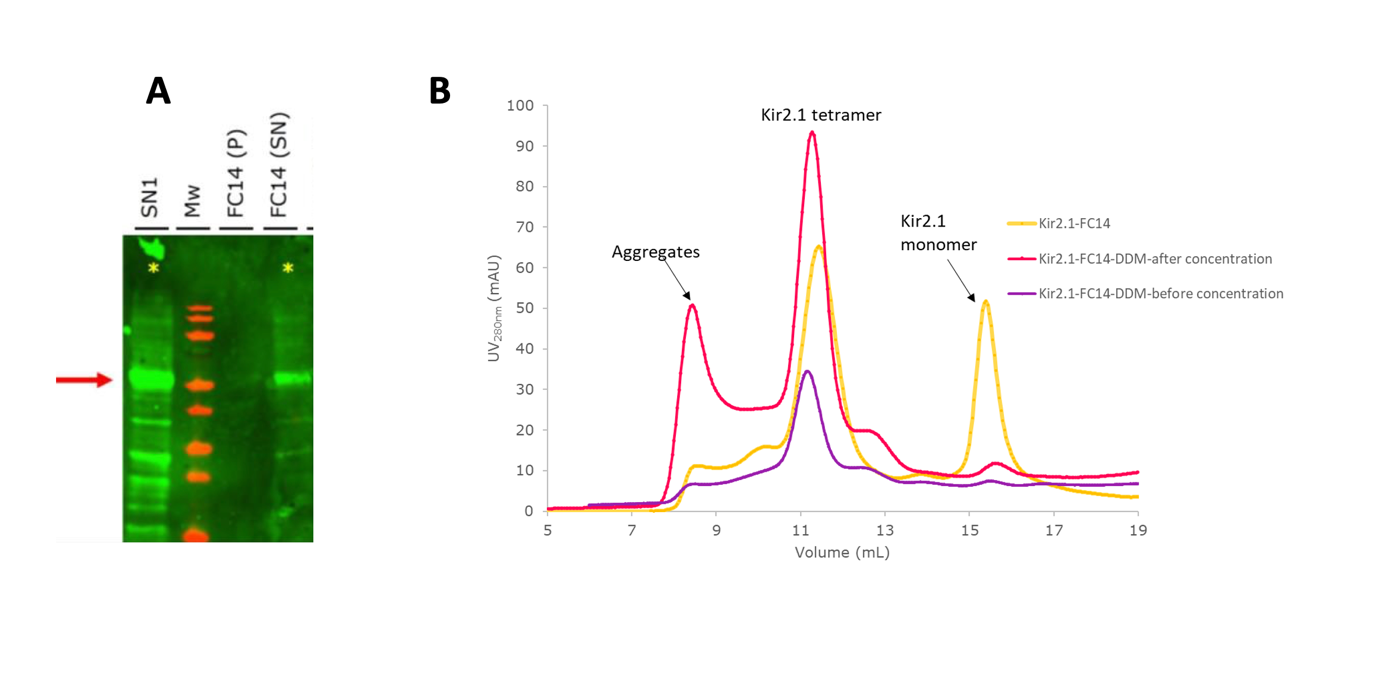

**Figure S3.** **Solubilization and purification of Kir2.1-WT using FC14 (tetradecyl-phosphocholine).** **A.** Western blot of membrane extraction of Kir2.1-WT using FC14. P indicates the unsolubilized pellet and SN the solubilized supernatant. **B.** SEC profile on Superdex® 200 (10/300) GL column (Cytiva) of Kir2.1-WT extracted in 0.5% FC14. Kir2.1 purified in 0.03% FC14 (yellow curve) elution presents two peaks; one corresponding to tetramers at 11.43 mL (R_H_ 60 Å) and another one corresponding to monomers at 15.43 mL (R_H_ 31.3Å). The purification after exchange to 0.03% DDM (purple curve) resulted in a single peak corresponding to tetramers (R_H_ = 63 Å). This sample tended to aggregate when concentrating by a factor of 25 (red curve).

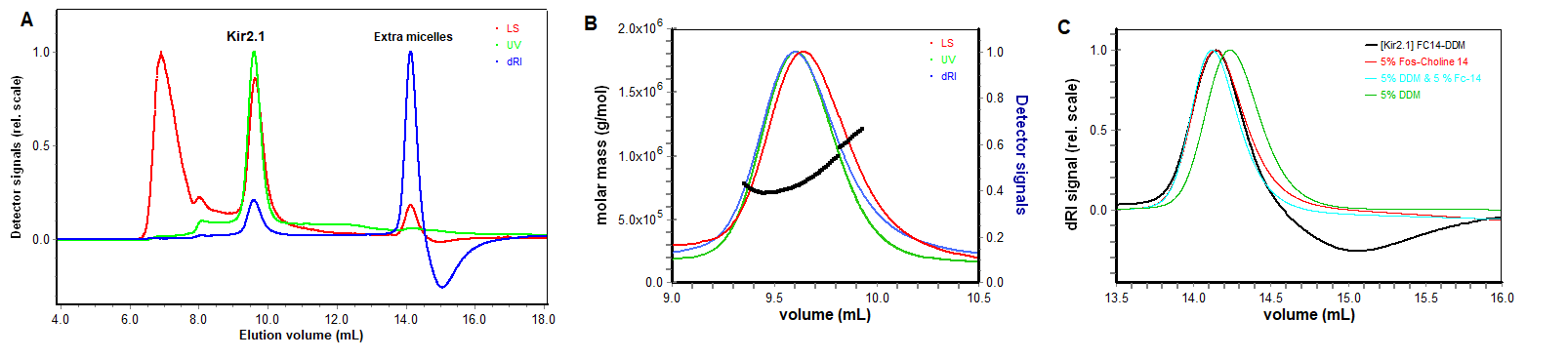

**Figure S4. SEC-MALLS-RI** **analysis of Kir2.1-FC14-DDM. A.** SEC-MALLS-RI profile of Kir2.1-WT FC14-DDM eluted on Superdex® 200 (10/300) GL column (Cytiva) with 0.03% DDM in SEC buffer (20 mM Tris-HCl pH 7.4, 150 mM KCl, 1 mM EDTA). The LS, UV, and ∆RI signals are shown in red, green, and blue, respectively; **B.** Zoom on the peak corresponding to tetramers showing no alignment of RI and UV signals with LS signal; **C.** Superposition of ∆RI signals of 5% DDM (green), 5% FC14 (red), 5% FC14-DDM mix (cyan) and extra micelles from Kir2.1-WT FC14-DDM (black). This shows that extra micelles from the Kir sample are not DDM micelles and that the exchange of FC14 by DDM on the affinity column is not complete.

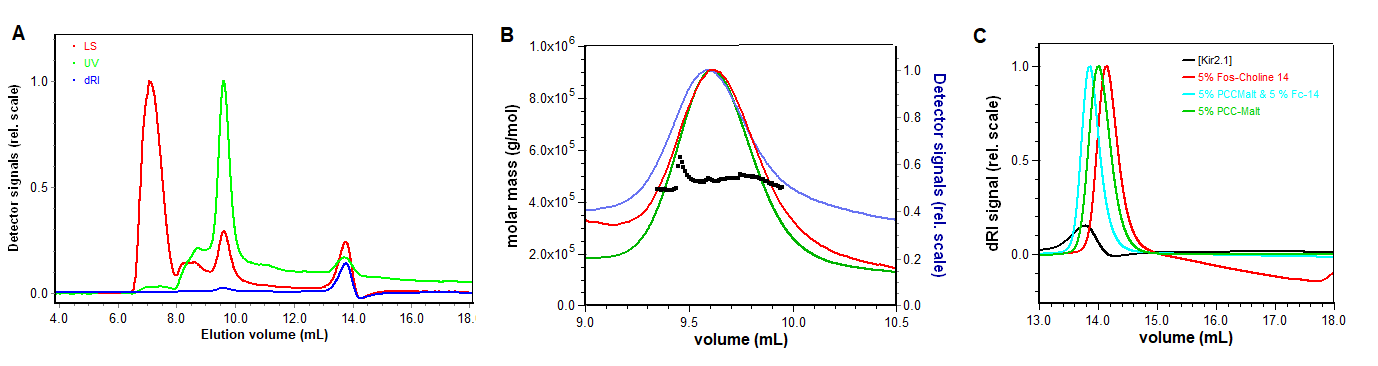

**Figure S5. SEC-MALLS-RI** **analysis of Kir2.1-FC14-PCC-MALT. A.** SEC-MALLS-RI profile of Kir2.1 FC14-PCC-MALT eluted on Superdex® 200 (10/300) GL column (Cytiva) with 0.03% PCC-Malt SEC buffer. The LS, UV and ∆RI are shown in red, green, and blue, respectively; B. Zoom on the peak corresponding to tetramers showing no alignment of UV, LS with RI signal, not allowing accurate calculation of molar mass fractions; **C.** Superposition of ∆RI signals for 5% PCC-Malt (cyan), 5% FC14 (red), and 5% FC14-PCC-Malt mix (green) and extra micelles from Kir2.1-WT-FC14-PCC Malt (black) eluted with 0.03% PCC-Malt SEC buffer. The extra micelles of the Kir sample (in black) eluted slightly before PCC-Malt micelles only.

**Methodology of mass calculation from SEC-MALLS experiments**

SEC-MALLS is a powerful technique to assess both the homogeneity and the molar mass of the biomolecules in solution. The main advantage of this method relies on the fact that no assumption, both on molecular conformation and molecular model, is necessary to determine the molar mass. In consequence, this method is very convenient to analyze purified membrane proteins solubilized in detergents. SEC-MALLS analysis permits the determination of the molar mass of the membrane protein complex, the amount of detergent bound onto the membrane protein, and the presence (or the excess) of the protein-free micelles.

The principle of this technique consists in the simultaneous difference measurement between the sample after SEC separation and the buffer (i.e. the mobile phase), the static light scattering (LS) at three different angles (∆I_θ_), the absorbance at 280 nm (∆A_280_) and the refractive index (∆RI).

Molar masses of each species separated by SEC are calculated using the protein conjugate mode in the Astra software (from Wyatt Technology) following successive defined steps.

First, the light scattering difference at each angle (∆I_θ_) is converted into Rayleigh ratio through the relationship:

$\Delta R\left( \theta\right)=\frac{\Delta I_{\theta}}{I_{t}}\cdot\left( \frac{n_{0}}{n_{t}} \right)^{2}\cdot R_{t}$	(1)

Where *n_0_*, *n_t_*, *I_t_*, and *R_t_* represent the refractive index of the mobile phase and toluene, the toluene scattering intensity at 90° and the toluene Rayleigh ratio at 90°, respectively.

The molar mass of the complex can then be obtained from the Rayleigh ratio:

$\Delta R\left( \theta\right)=K\cdot M_{wcomplex}\cdot\left[ complex \right]\cdot\left( \frac{ⅆn}{ⅆc} \right)_{complex}^{2}\cdot P\left( \theta\right)\cdot S\left( \theta\right)$	(2)

Where *K*, *Mw*, $\left( \frac{ⅆn}{ⅆc} \right)_{complex}$, *P*(θ), *S*(θ) are an optical constant, the weight-average molecular mass of the complex, the increment refractive index of the complex, the form factor of the complex, and the structure factor (formally equal to 1 in chromatographic condition), respectively.

Second, the signals from both UV and RI detectors are used to determine the concentration of the complex. Using both absorbance and refractive index, the concentration of the complex can be written as follows:

$\left[ complex \right]=\frac{\Delta A_{280}}{\varepsilon_{complex}}=\frac{\Delta RI}{\left( \frac{ⅆn}{ⅆc} \right)_{complex}}$	(3)

Using the detergent-to-protein weight ratio (δ), we obtain the following equation:

$\left[ complex \right]=\frac{\Delta A_{280}}{\left( \frac{1}{1+\delta} \right)\cdot A_{280p}+\left( \frac{\delta}{1+\delta} \right)\cdot A_{280d}}=\frac{\Delta RI}{\left( \frac{1}{1+\delta} \right)\cdot\left( \frac{ⅆn}{ⅆc} \right)_{p}+\left( \frac{\delta}{1+\delta} \right)\cdot\left( \frac{ⅆn}{ⅆc} \right)_{d}}$	(4)

In which $A_{280p}$, $A_{280d}$, $\left( \frac{ⅆn}{ⅆc} \right)_{p}$, $\left( \frac{ⅆn}{ⅆc} \right)_{d}$ are the extinction coefficient of the protein and detergent, the refractive index increment of the protein and detergent, respectively. Then, from equation (4), the Protein conjugate mode allows one to determine the value of δ.

In the third step, the inverse of the weight-averaged molar mass of the complex is determined from the intercept of the Zimm plot:

$\frac{K\cdot\left[ complex \right]\cdot\left( \frac{ⅆn}{ⅆc} \right)_{complex}^{2}}{\Delta R\left( \theta\right)}=\frac{1}{M_{wcomplex}\cdot P\left( \theta\right)}$	(5)

Finally, the weight-averaged molar mass of the protein and bound detergent are given by:

$Mwprotein=\frac{M_{wcomplex}}{1+\delta}$	(6)

When the complex is monodisperse, the normalized signals from all detectors are superimposed. Under these conditions, the weight-averaged molar mass of the complex is equal to the average molar mass.

Before doing a SEC-MALLS experiment on a complex, a template must be first done with a monodisperse standard protein (BSA for instance) in the running buffer. Following this run, three steps must be done:

- the normalization of the LS detectors to the LS detector at 90°, in order to correct the scattering volume difference;

- the inter-detector delay correction, to take into account the volume between each detector;

- the band broadening correction to take into account the sample diffusion through both the detectors and the inter-detector tubing.

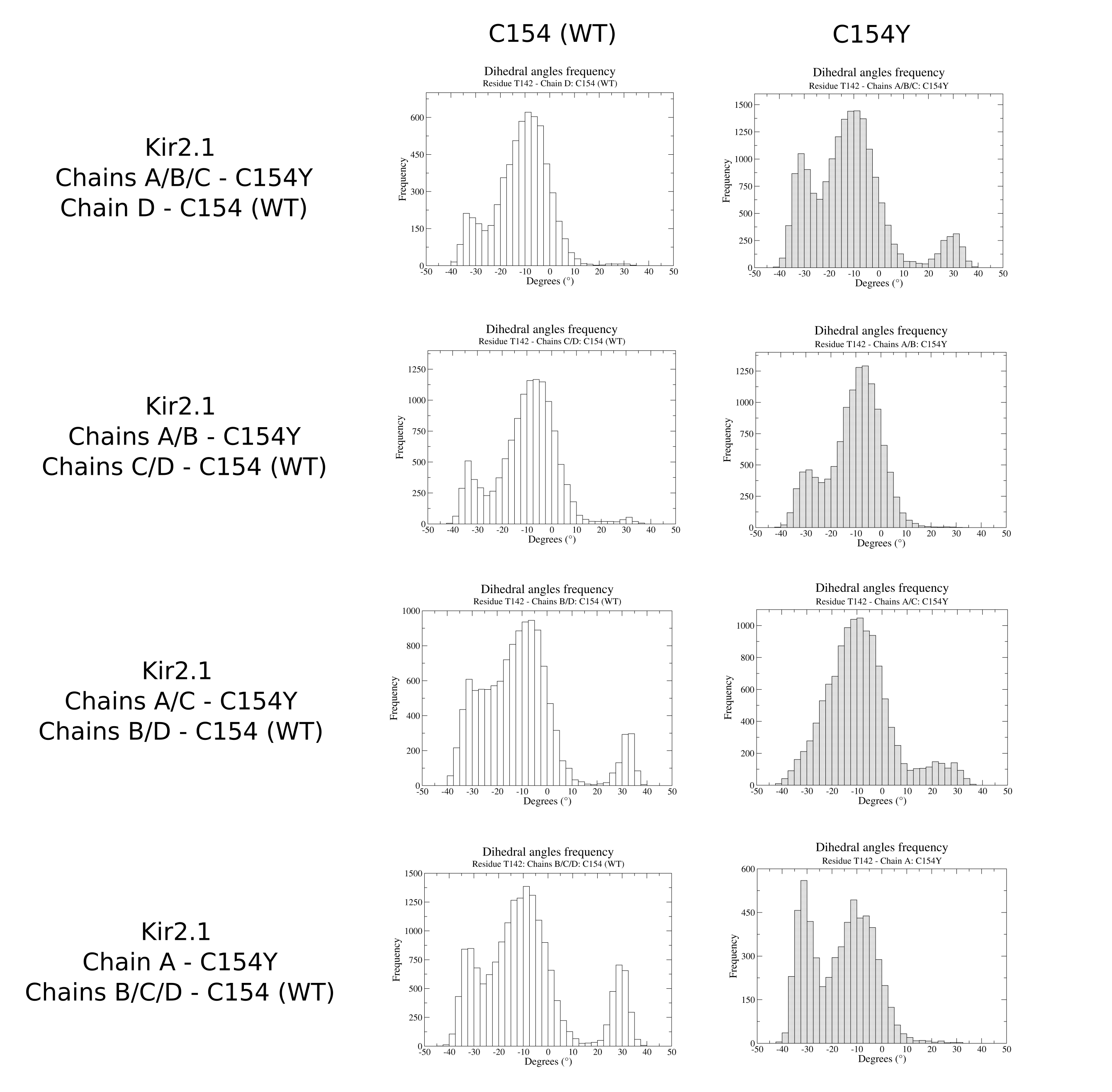

**Figure S6.** Frequency of dihedral angles for the T142 residue obtained along the 200 ns of MD simulations in triplicate of the Kir2.1 channel cryo-EM structure after incorporating the C154Y mutation in three subunits of the tetramer (Chains ABC_C154Y_, Chain D_WT_), in two adjacent (side-by-side) subunits (Chains AB_C154Y_, Chains CD_WT_), in two diagonally opposite subunits (Chains AC_C154Y_, Chains BD_WT_), and in one subunit (Chain A_C154Y_, Chains BCD_WT_). No values below −50° or above 50° were identified.

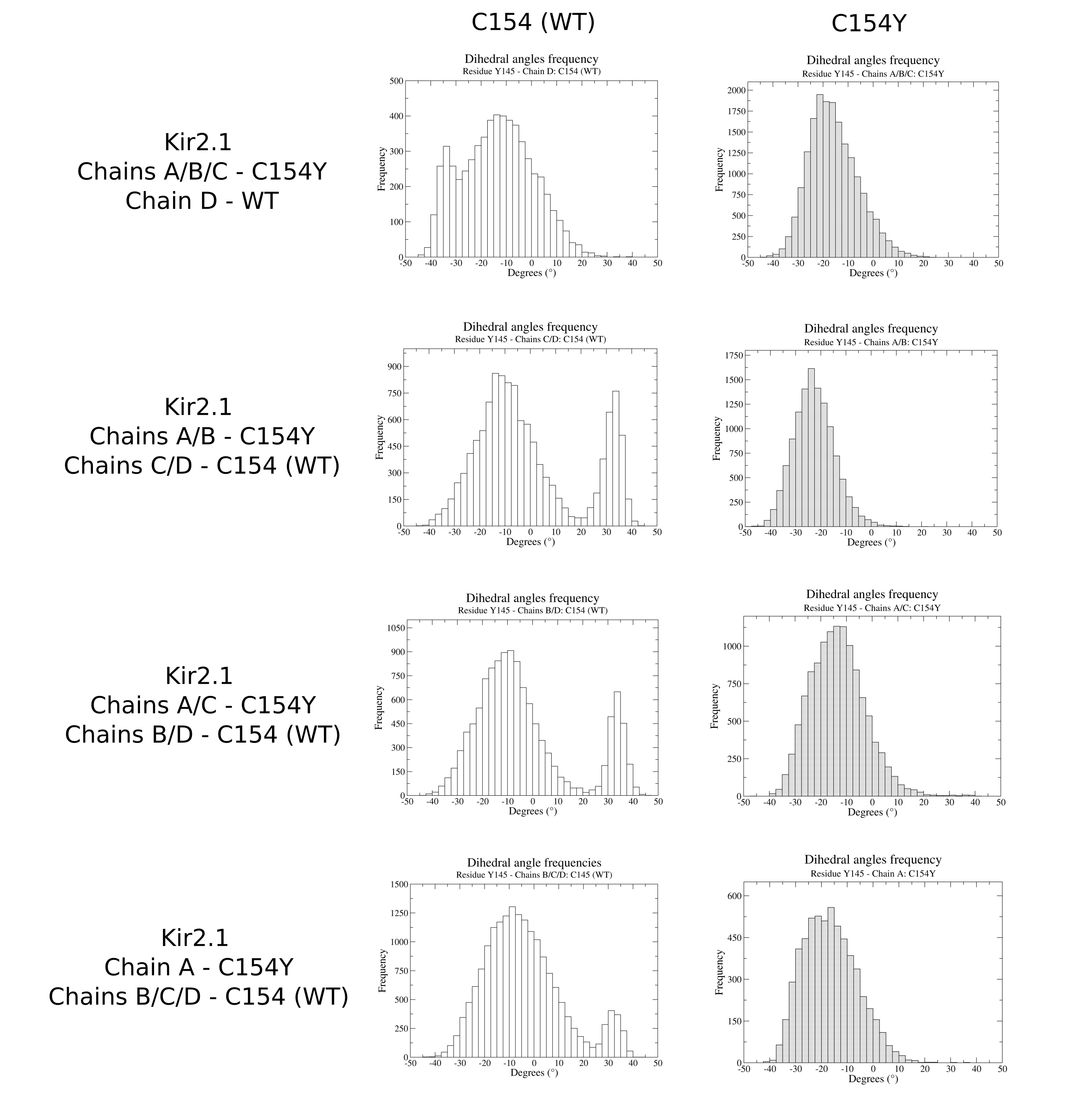

**Figure S7.** Frequency of dihedral angles for the Y145 residue obtained along the 200 ns of MD simulations in triplicate of the Kir2.1 channel cryo-EM structure after incorporating the C154Y mutation in three subunits of the tetramer (Chains ABC_C154Y_, Chain D_WT_), in two adjacent (side-by-side) subunits (Chains AB_C154Y_, Chains CD_WT_), in two diagonally opposite subunits (Chains AC_C154Y_, Chains BD_WT_), and in one subunit (Chain A_C154Y_, Chains BCD_WT_). No values below −50° or above 50° were identified.

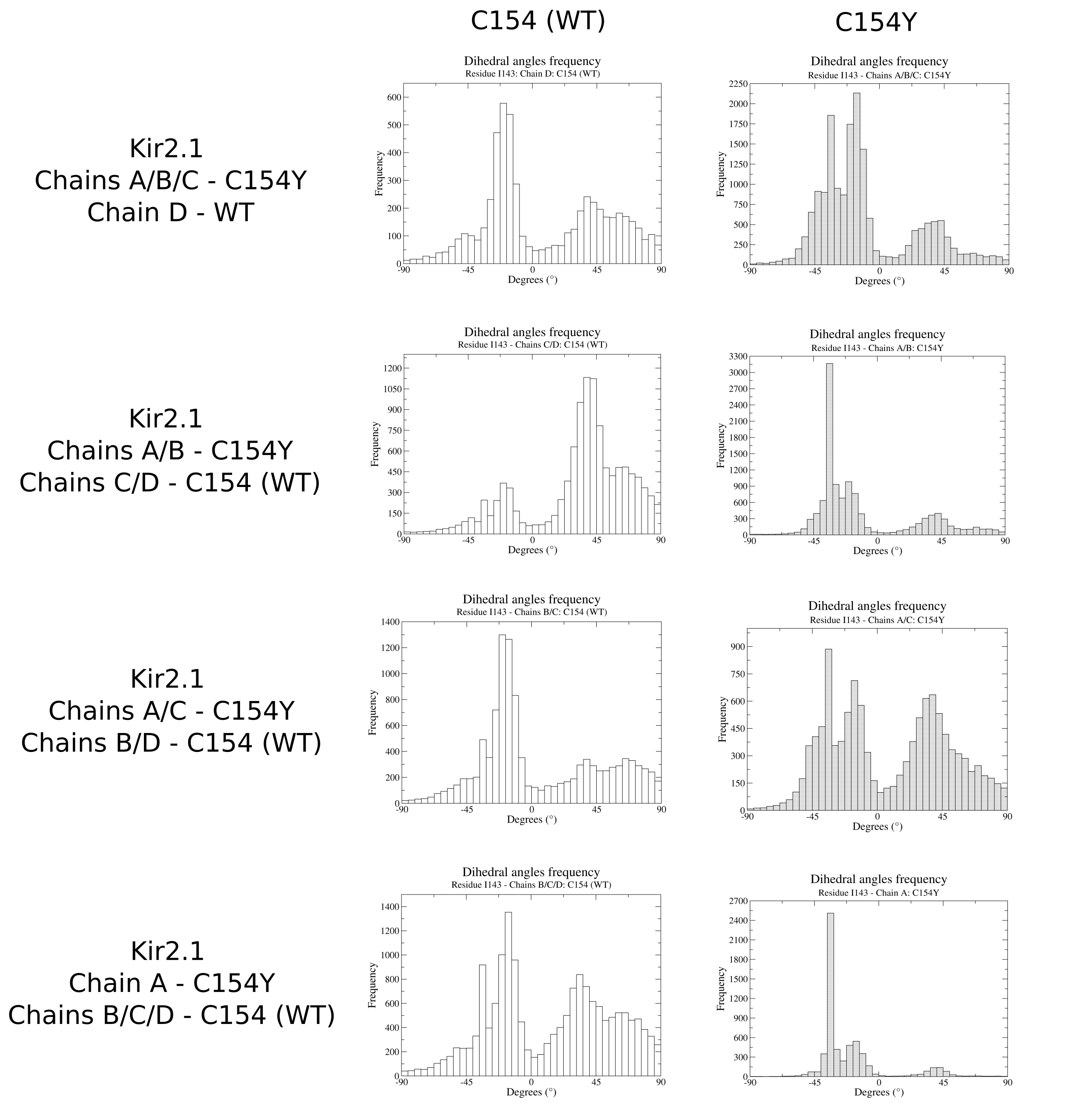

**Figure S8.** Frequency of dihedral angles for the I143 residue obtained along the 200 ns of MD simulations in triplicate of the Kir2.1 channel cryo-EM structure after incorporating the C154Y mutation in three subunits of the tetramer (Chains ABC_C154Y_, Chain D_WT_), in two adjacent (side-by-side) subunits (Chains AB_C154Y_, Chains CD_WT_), in two diagonally opposite subunits (Chains AC_C154Y_, Chains BD_WT_), and in one subunit (Chain A_C154Y_, Chains BCD_WT_). The values below −90° or above 90° are negligible.

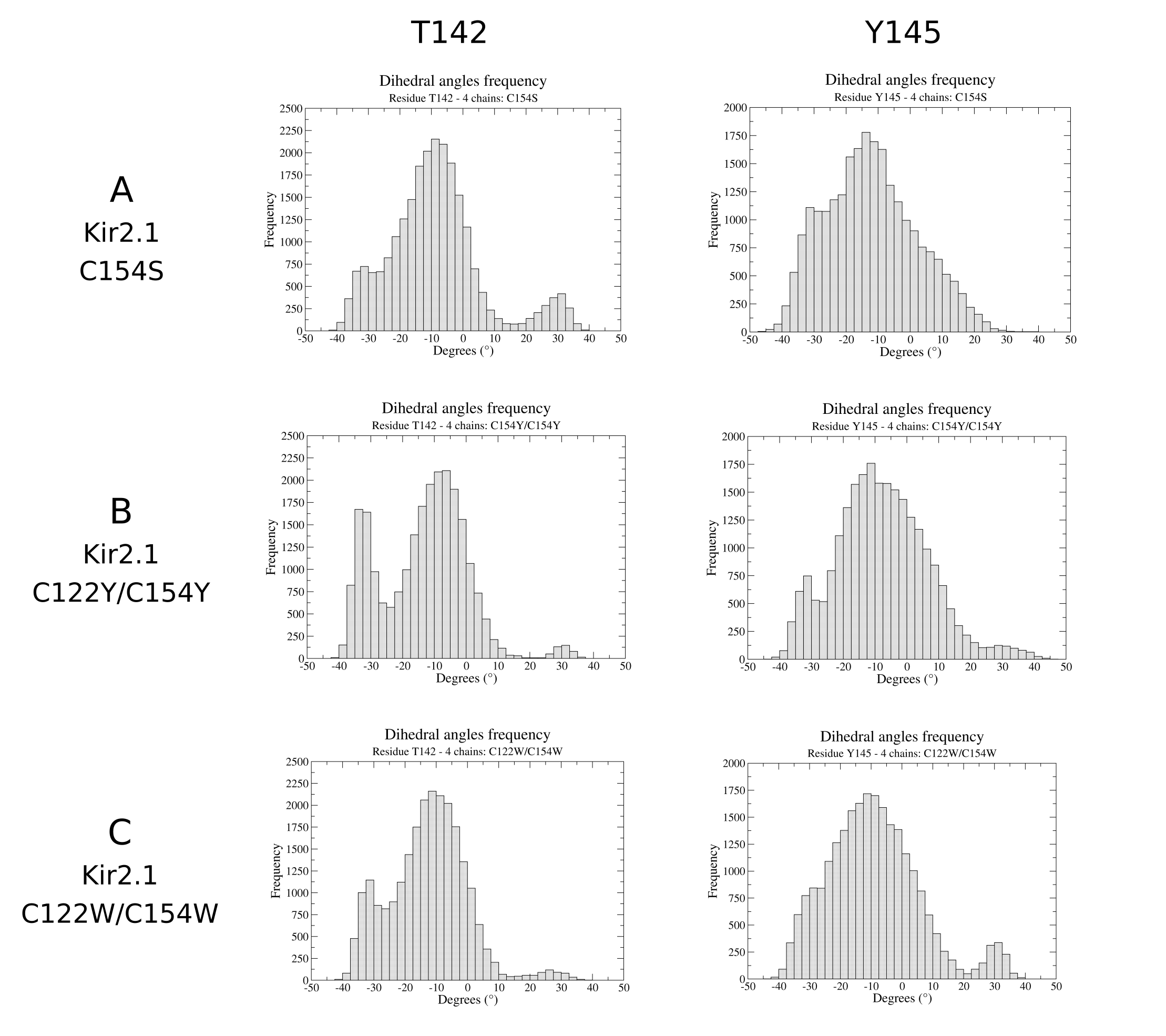

**Figure S9.** Frequency of dihedral angles for the T142 (left column) and Y145 (right column) residues obtained along the 200 ns of MD simulations (one replica) of the Kir2.1 channel cryo-EM structure after incorporating the C154S (panel **A**), C122Y/C154Y (panel **B**) and C122W/C154W (panel **C**) mutations in the four subunits of the tetramer.

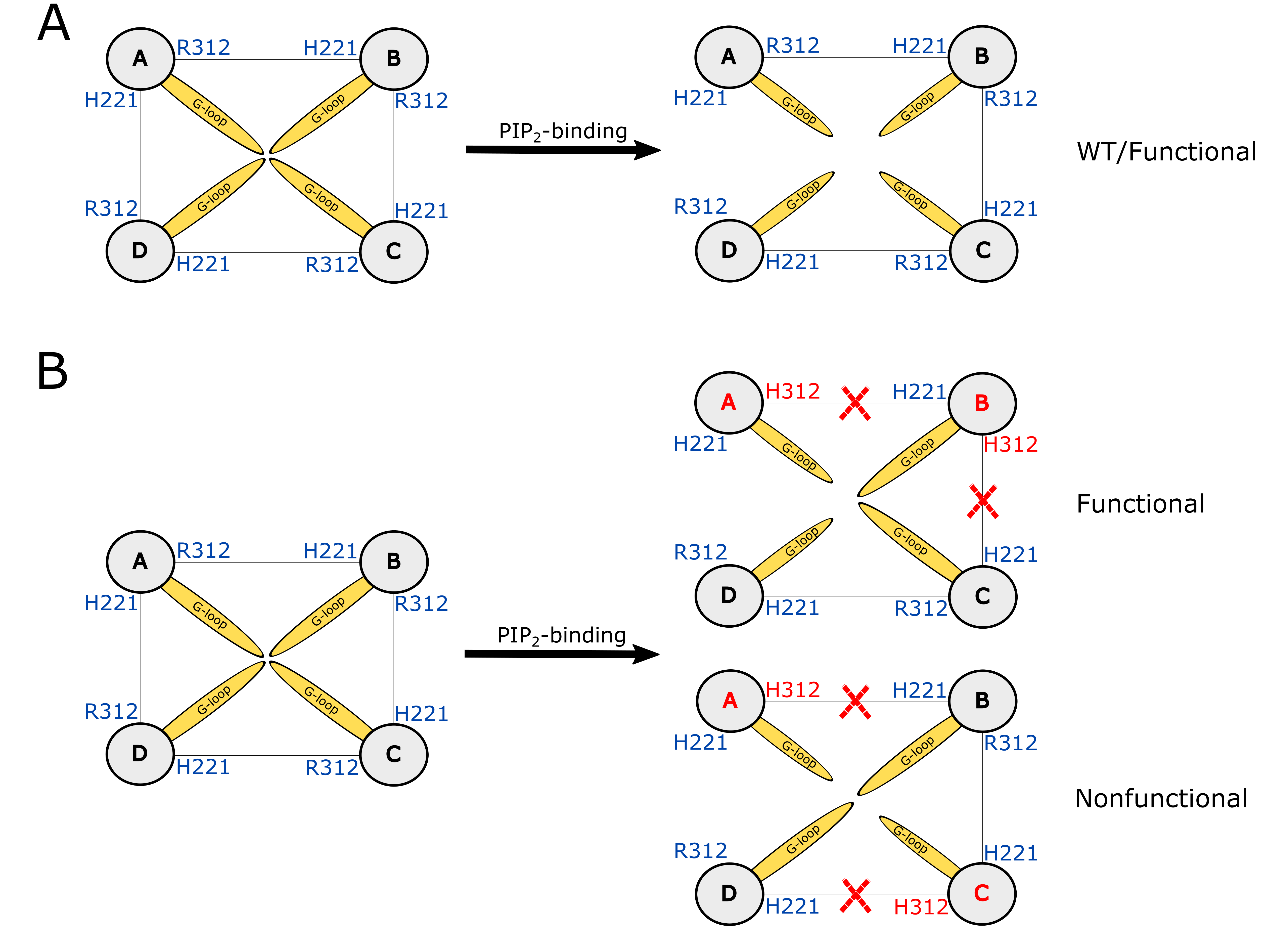

**Figure S10. Structural hypothesis on the loss of function of Kir2.1-R312H mutant when the mutation is placed in two of the four subunits of the tetramer. A.** Schematic representation of the opening of the G-loop induced by the PIP_2_-binding in the Kir2.1-WT (functional channel). **B.** Schematic representation of the opening of the G-loop induced by the PIP_2_-binding in the Kir2.1-R312H mutant when the R312H mutation is placed on side-by-side subunits (top) or diagonally opposed subunits (bottom). When the R312H mutation is placed in two diagonally opposite subunits, the G-loops of the R312H-containing subunits close the center of channel pore, resulting in a nonfunctional channel. In contrast, when the R312H mutation is placed on side-by-side subunits, part of the channel pore remains unblocked, resulting in a functional channel.

**Table S1.** Root mean square fluctuation (rmsf) values calculated along the 200 ns of MD simulations in triplicate of the Kir2.1 channel cryo-EM structure after incorporating the C154S, C122Y/C154Y, and C122W/C154W mutations in the four subunits of the tetramer.

|  | **Kir2.1**  **Chains A/B/C/D: C154S** | **Kir2.1**  **Chains A/B/C/D: C122Y/C154Y** | **Kir2.1**  **Chains A/B/C/D: C122W/C154W** |
| --- | --- | --- | --- |
| **Position 154** | 1.80±0.25 | 1.82±0.18 | 1.68±0.06 |
| **Loop 147-153** | 1.64±0.19 | 1.77±0.14 | 1.60±0.08 |
| **Selectivity filter (142-146)** | 1.27±0.18 | 1.35±0.12 | 1.22±0.05 |

.

Table S2. Cryo-EM data collection and refinement statistics

| Data Collection |  |  |
| --- | --- | --- |
| Microscope |  | Titan Krios |
| Detector |  | Falcon 4 |
| Voltage (kV) |  | 300 |
| Pixel size (Å/pixel) |  | 0.73 |
| Electron dose per image |  | 40 |
| Exposure time (s) |  | 2.35 |
| Defocus range (μm) |  | −1.2 to −2.4 |
| Movies collected |  | 10 762 |
| Movies used |  | 10 067 |
| Reconstruction |  |  |
| Particles used for final reconstruction |  | 139 500 |
| Final Resolution (Å) (FSC = 0.143) |  | 6.03 |
| Model composition |  |  |
| Model coverage |  | 4 chains, each chain in the range 46-60 and 70-362 |
| Non-hydrogen atoms |  | 9864 |
| Refinement |  |  |
| CC (calculated by Chimera) |  | 0.86 |
| CC (mask) |  | 0.65 |
| CC (volume) |  | 0.63 |
| CC (peaks) |  | 0.34 |
| d_model_ (Å) |  | 6.00 |
| r.m.s deviations |  |  |
| Bond length (Å) |  | 0.003 |
| Bond angles (°) |  | 0.856 |
| Validation |  |  |
| MolProbity Score |  | 2.81 |
| Clash Score |  | 30.34 |
| Ramachandran Plot Outliers (%) |  | 0.99 |
| Rotamer outliers (%) |  | 0 |
| Cβ outliers (%) |  | 0 |
| EMDB ID |  | EMD-18595 |
| PDB ID |  | 8QQL |
